# Adolescent drinking causes a loss of aspartoacylase-expressing oligodendrocytes and hypomyelination of anterior cingulate and corpus callosum axons in male mice, but not females

**DOI:** 10.64898/2026.04.01.715654

**Authors:** Said Akli, Annabelle Flores-Bonilla, Sirisha Nouduri, Samuel Scott, Heather N. Richardson

## Abstract

Adolescent binge drinking is a strong predictor of alcohol use disorder and related mental health outcomes in adulthood, which may be due to disruptions in myelination during this dynamic period of brain development. White matter expansion in frontal regions during adolescence is essential for mature decision-making and stress regulation, yet the cellular mechanisms by which alcohol disrupts this process remain poorly understood. We used multi-label immunofluorescence and confocal microscopy to visualize proteins in oligodendrocyte lineage cells and myelin ensheathment of axons in the anterior cingulate cortex (Cg1) and corpus callosum (CC) following four weeks of episodic voluntary binge drinking using the Drinking-in-the-Dark model in adolescent male and female C57BL/6NJ mice beginning on postnatal day 28. Contrary to our initial hypothesis that alcohol targets early-stage oligodendrocyte precursor cells (OPCs), binge drinking selectively depleted mature oligodendrocytes expressing aspartoacylase (ASPA) in the Cg1 and CC of male mice, but not females. This enzyme is essential for lipid biosynthesis and myelin production, and this cell-specific loss was accompanied by significant hypomyelination of axons only in males. These findings identify a later maturational stage of oligodendroglial development as a sex-dependent target of alcohol, advancing our mechanistic understanding of prefrontal myelin deficits in adolescent drinking. Furthermore, ASPA emerges as a potential therapeutic target for alcohol use disorder and demyelinating diseases, with differential vulnerability across sex carrying important implications for adult neurodevelopmental outcomes.

## Introduction

Over 134 million teenagers and adults consumed alcohol in the United States in 2024, with 43.1% reporting binge drinking within the past month (Substance Abuse and Mental Health Services Administration, 2025a). While the overall prevalence of binge drinking among adult men and women declined from 2023 to 2024, rates among adolescents aged 12-17 has remained unchanged over the last few years (2.9% in young males and 4.1% in young females, (Substance Abuse and Mental Health Services Administration, 2025b, 2025c)). Binge drinking is the rapid consumption of alcohol that produces blood alcohol concentrations of 0.08 g/dL or higher within two hours, and this is the most common, costly, and deadly type of excessive drinking (National Institute on Alcohol Abuse and Alcoholism, 2004; Stahre et al., 2014; Sacks et al., 2015). Binge drinking during adolescence is strongly associated with the later development of alcohol use disorder (AUD), a chronic relapsing condition characterized by maladaptive alcohol use that persists despite physiological and psychological distress (Chou and Pickering, 1992; American Psychiatric Association, 2013; Addolorato et al., 2018). The neurocircuits implicated in AUD and co-occurring mental health conditions are known to differ with sex (Flores-Bonilla and Richardson, 2020; Lees et al., 2020; Radke et al., 2021; Bowen et al., 2022). Determining how adolescent drinking affects maturation of these neurocircuits in males and females may help elucidate the underlying causes of cognitive and behavioral health outcomes in adulthood.

Brain maturation during adolescence includes the expansion of myelinated white matter fiber tracks in frontal brain regions, which is thought to contribute to improved decision-making, cognition, stress-regulation, and impulse control in adulthood (Gogtay et al., 2004). Oligodendrocytes (OLs) are the glial cells that form insulating myelin sheaths by wrapping concentric layers of lipid-rich processes around axons (Simons and Nave, 2016; Stadelmann et al., 2019)

Myelin sheaths restrict membrane conductance to the nodes of Ranvier that are enriched with voltage-gated ion channels, leading to faster neurotransmission through saltatory conduction (Seidl, 2014). The functional benefit of prefrontal myelination during adolescence has been demonstrated in preclinical animal studies. For example, myelination of axons extending from the corpus callosum into the anterior cingulate cortex early in adolescence (McDougall et al., 2018; Drzewiecki et al., 2020) corresponds with a marked increase in the transmission speed of action potentials along these axons in male rats (McDougall et al., 2018).

In addition to the increased risk of AUD, adolescent drinking has also been linked to other negative health outcomes in adulthood including impaired higher executive functions, enhanced reward and sensation-seeking, and a dysregulated response to stress (McCarty et al., 2004; Peters et al., 2015; Squeglia et al., 2015; Elsayed et al., 2018), reviewed in (Crews et al., 2016).These functional deficits observed in adulthood could be due to alcohol-induced disruptions in myelination during adolescent development, as reviewed in (Rice and Gu, 2019). Indeed, lower expression of myelin-associated genes and myelin deficits have been observed in the anterior cingulate following adolescent drinking in humans, rats and mice, with greater effects in males compared to females (Vargas et al., 2014; Wolstenholme et al., 2017; Morris et al., 2019; Tavares et al., 2019).

Establishing a causal link between alcohol-induced myelin loss in adolescence and negative health outcomes in adulthood requires a deeper understanding of the cellular mechanisms underlying these changes in myelin, yet these remain elusive. Alcohol may be disrupting axonal myelination in the developing brain of adolescents by targeting OL precursor cells (OPCs) and triggering apoptotic cell death or preventing these cells from differentiating into OLs. In support of this, there is evidence of OPC apoptosis and attenuated differentiation in human fetal brain tissue following alcohol exposure during embryonic development (Darbinian et al., 2021). Similar results have also been reported in rodents exposed to prenatal alcohol (Newville et al., 2017). Alternatively, adolescent alcohol may be targeting this lineage after differentiation, as has been observed in fetal brains of macaques following alcohol exposure (Creeley et al., 2013).

The goal of the current study was to identify cellular mechanisms that could account for myelin changes after repeated binge drinking in adolescence. Using multi-label immunofluorescence and confocal microscopy we tracked and quantified cellular proteins in oligodendroglia lineage cells and myelin ensheathment of axons in the anterior cingulate cortex and corpus callosum following repeated cycles of voluntary binge drinking of alcohol (or water) using the Drinking-in-the-Dark model in adolescent male and female mice. Contrary to our initial hypothesis, we found evidence of oligodendroglia lineage disruption in later stages of cellular development when mature OLs are making the aspartoacylase enzyme necessary for lipid biosynthesis and myelin production. Most notably, these cells were especially sensitive to high levels of alcohol, but the effects were unique to males—explaining why reduced myelin density was not observed in binge drinking females. These findings offer insight into the mechanisms underlying prefrontal myelin deficits associated with adolescent alcohol use, and differential sensitivity across sex may have important implications for adult outcomes.

## Materials and Methods

### Animals

C57BL/6NJ wildtype male and female mice arrived at 3 weeks of age from Jackson Laboratory (Bar Harbor, ME; stock #005304) and were housed in same sex groups of 3-5 mice per cage. All care of and experimental procedures with animals were performed in strict compliance with the University of Massachusetts Amherst Institutional Animal Care and Use Committee and the NIH Guide for the Care and Use of Laboratory Animals.

### Drinking-in-the-Dark (DID)

Male and female C57BL/6NJ wild-type mice (n=8/alcohol group/sex) were exposed to voluntary alcohol binge drinking using the well-established limited access Drinking-in-the-Dark (DID) procedure (Rhodes et al., 2005a; Thiele and Navarro, 2014; Thiele et al., 2014; Wilcox et al., 2014). Mice were transferred to a reverse cycled room (11:00 lights OFF/ 23:00 lights ON) for acclimatization for 1 week and were single housed at postnatal day (PD) 28 for DID. DID began 3 hours into the dark cycle on PD28. Mice were given access to either 20% (v/v/) alcohol (or water for controls) for 2 hours on 3 consecutive days, followed by a “binge day” with 4 hours of access. This was followed by a 3-day rest period in the home cage without access to alcohol before the next drinking DID cycle began. Mice completed four DID cycles in total from PD28-56. To access drinking levels, bottles were weighed before and after each bout to provide a measure used to calculate daily g/kg intake of alcohol for each animal or ml/kg water intake for control animals. Spillage bottles containing either water or alcohol were handled identically to the experimental bottles to estimate the loss of liquid during the manipulation and due to evaporation. The average spillage volumes were subtracted from the experimental volumes of alcohol or water. Negative values were counted as 0 g/kg intake for alcohol or water.

### Intracardial perfusions and brain tissue processing

Four days after the last drinking bout (PD56), animals were deeply anesthetized with pentobarbital sodium solution (Nembutal, 200 mg/kg *ip*). They were intracardially perfused with 0.9% saline (room temperature) for 5 min, followed by 4% paraformaldehyde / 0.1M sodium tetraborate (chilled to 4°C, pH 9.4) for 5 min at a rate of 6 ml/min. Brains were rapidly extracted and post-fixed overnight in 4% paraformaldehyde at 4°C. Brains were then immersed in a 10% sucrose/phosphate buffered saline solution at 4°C for 1 day and 30% sucrose/phosphate buffered saline solution for another 24 hours. Brains were then snap frozen by submerging briefly in -50°C isopentane (2-methylbutane; Thermo Fisher Scientific, catalogue #03551-4, Fair Lawn, NJ, USA) and stored at -80°C until cryo-sectioning. A freezing sliding microtome was used to slice 35 μm thick coronal serial sections that were collected in strict anatomical order and stored in cryoprotectant solution in a 1:6 series at -20°C until immunofluorescence experiments were performed.

### Immunofluorescence

In Experiment 1, we used free-floating sections at AP+1.7mm to align anatomically with previous studies showing prefrontal myelin loss after adolescent exposure to alcohol drinking in rats or binge intragastric alcohol administration in mice (Vargas et al., 2014; Papp-Peka et al., 2016; Rice et al., 2019; Tavares et al., 2019). In Experiment 2, we expanded the sampling region to include more sections (AP+2.0mm, AP+1.7mm, AP+0.9mm) and test whether oligodendroglial cellular changes extend beyond the medial prefrontal cortex. For both experiments, floating sections were first thoroughly washed with PBS to remove the cryoprotectant storage solution. This was followed by washes in 0.3% Triton-X in PBS (PBS-Tx) to enable primary antibodies to permeate cytoplasmic and nuclear membranes and bind to target proteins. Tissue samples were incubated in 50 mM ammonium chloride solution (NH_4_Cl/PBS-Tx) for 30 minutes at 25°C to quench autofluorescence followed by rinses in PBS-Tx. They were then incubated in a 3% hydrogen peroxide/PBS solution for 30 minutes at 25°C to block endogenous peroxidase. To reduce non-specific binding of secondary antibodies, tissues were incubated in a 5% normal horse serum/PBS-Tx solution for 1 hour at 25°C. Samples were then thoroughly washed in PBS-Tx, followed by incubation with primary antibodies specific to each experiment overnight at 4°C (see **Table 1** for dilutions). After several washes in PBS-Tx, sections were incubated in secondary antibodies at room temperature (25°C) for 2 hours. Sections were then washed in PBS and incubated in a Cy3-conjugated streptavidin/PBS-Tx solution for 1 hour at 25°C to amplify and visualize the CC1 signal. Following PBS rinses, nuclei were then fluorescently stained using 0.4 mg/ml diamidinophenolindole (DAPI), which binds to the adenine–thymine-rich regions in DNA (Kapuscinski, 1995). Sections were washed in PBS and mounted on subbed glass slides and allowed to thoroughly dry in a dark room for a minimum of 12 hours. Slides were next cleared in xylenes and DPX mounting media was used to adhere glass coverslips (Fisher Scientific #50980370).

**Table 1.** Antibodies and amplification reagents used for immunohistochemical experiments.

| <u>Reagent</u> | <u>Species</u> | <u>Supplier</u> | <u>Cat #</u> | <u>Dilution</u> |
| --- | --- | --- | --- | --- |
| **Primary antibody** |  |  |  |  |
| MOG | Rabbit | Abcam | Ab32760 | 1:1000 |
| PDGFRα+ | Goat | R&D System | AF1062 | 1:100 |
| QKI-7 (clone-CC1) | Mouse | Millipore | MABC200 | 1:400 |
| ASPA | Rabbit | Millipore | ABN1698 | 1:1000 |
| **Secondary antibody** |  |  |  |  |
| Anti-Rabbit (AlexaFluor 647) | Donkey | Jackson ImmunoResearch |  | 1:500 |
| Anti-Goat (AlexaFluor 588) | Donkey | Jackson ImmunoResearch |  | 1:500 |
| Biotinylated a Mouse (IgG) | Horse | Vector Laboratories | BA-2000 | 1:200 |
| Cy3-Streptavidin |  | Jackson ImmunoResearch | 016-160-084 | 1:2000 |

### *Experiment 1:* Test for evidence of reduced oligodendrocyte progenitors and mature cells accompanying myelin loss following adolescent drinking

We used multi-label immunofluorescence to detect and visualize oligodendroglia- and myelin-related proteins in medial prefrontal white and gray matter regions at the location where axons of the corpus callosum forceps minor (CC_FM_) branch out into the cortical layers of the anterior cingulate (Cg1). This region of interest was selected based on previous reports identifying it as a sight of dynamic adolescent myelination (McDougall et al., 2018; Drzewiecki et al., 2020) and sensitivity to alcohol (Vargas et al., 2014; Tavares et al., 2019). Antibodies and reagents are detailed in **Table 1**. To assess myelinated fiber density, we used a primary antibody that recognizes myelin oligodendrocyte glycoprotein (MOG), which is located on the outer layer of the myelin sheath (Clements et al., 2003; Ambrosius et al., 2020). Primary antibodies that recognize proteins that are expressed at specific oligodendroglia lineage stages were used to delineate between oligodendroglia precursor cells (OPCs) and oligodendroglia (OLs) for cellular density analyses: anti-platelet-derived growth factor receptor alpha (PDGFRɑ) primary antibody for OPCs and anti-CC1 primary antibody for OLs that express RNA-binding protein Quaking Isoform 7 (QKI-7). Fluorophore-conjugated secondary antibodies used to visualize these proteins were donkey anti-rabbit-AF647 for MOG, a donkey anti-goat-AF488 for PDGFRɑ, and a horse anti-mouse biotin antibody followed by incubation with Cy3-streptavidin for QKI-7.

### *Experiment 2:* Test for evidence of reduced aspartoacylase production in mature oligodendrocytes following adolescent drinking

To study the late maturational stages of cellular development we used a combination of antibodies to test for the presence or absence of the aspartoacylase enzyme in mature OLs (antibodies and reagents are summarized in **Table 1**). These analyses expanded the sampling region to include cingulum bundle (CC_cing_) axons, which extend into the Cg1 more posteriorly after crossing the midline. Mature OLs expressing QKI-7 were immunolabeled using anti-CC1 antibody and co-labeled with antibody against the aspartoacylase (ASPA) protein to determine if these mature OLs were producing the enzyme required to form and maintain myelin sheaths (Madhavarao et al., 2004; Francis et al., 2012, 2016; Grønbæk-Thygesen and Hartmann-Petersen, 2024; Takeda et al., 2024). This allowed us to delineate the specific maturational stage at which OLs were impacted by alcohol. In a subset of mice, sections were also immunolabeled for myelin basic protein to confirm ASPA-expressing cells were producing myelin sheath proteins. We therefore included incubations in chicken anti-MBP primary antibody and Alexa 488-conjugated goat anti-chicken antibody steps for those sections. Fluorophore-conjugated secondary antibodies used for detection were a horse anti-mouse biotin antibody followed by incubation with Cy3-streptavidin for QKI-7 and a donkey anti-rabbit-AF647 for ASPA.

#### Confocal microscopic imaging

All images in Experiment 1 were acquired on the A1R-TIRF confocal microscope and all images in Experiment 2 were acquired on the CREST-V2 confocal microscope at the UMass IALS Nikon Center for Excellence Light Microscopy Facility Core. Both microscopes are connected to the Nikon NIS-Elements platform to process and analyze images. To assess white matter, we sampled from the location where corpus callosum axons extend medially out into the layer VI of the anterior cingulate cortex. At the level of the medial prefrontal cortex this is called the forceps minor or CC_FM_ and at the level of the bed nucleus of the stria terminalis this is called the cingulum or CC_cing_. To assess prefrontal gray matter, we sampled from layers II/III within the anterior cingulate cortex (Cg1). In the first study, sections containing the medial prefrontal cortex (AP+1.7 mm distance from bregma) were triple immunofluorescent labeled for MOG, PDGFRɑ, and QKI-7, and counterstained with DAPI fluorescent nuclear stain. In the second study, sections were selected from the following anatomical locations: AP+2.0mm, AP+1.7mm, and AP+0.9mm distance from bregma. These sections were double immunofluorescent labeled for QKI-7 and ASPA, and counterstained with DAPI nuclear stain. Within each A/P distance from bregma and each hemisphere, one white matter (CC_FM_) and one gray matter (Cg1) region of interest (ROI) were selected for imaging. A 20X objective was used for acquiring z-stacks images for both the A1R-TIRF (Experiment 1) and CREST-V2 (Experiment 2) confocal microscopes.

#### Quantification of myelinated axons

Sections that were AP+1.7mm were used for analysis of myelin density in the ROIs. eneral Analysis 3 with NIS-AR was used to quantify MOG positive fiber density by thresholding the images in the infra-red channel (647nm). The percentage of area covered by myelinated fibers over the total area was calculated to quantify myelinated fiber density.

#### Quantification of oligodendroglial lineage cell populations

An optical configuration was designed within the NIS-Advanced Research (NIS-AR) software, allowing all imaging to occur with identical laser power and gain parameters. First, a 10x-stitched large image of the entire tissue section was acquired on the 405 nm channel for DAPI visualization of neural architecture. This image was used to digitally select our ROIs in each hemisphere. For each ROI, a 20x z-stack with a depth of 10µm and a step size of 1µm was acquired. A custom-designed semi-automatic cell counting program in General Analysis 3 within NIS-AR quantified multiple cell types in three-dimensions simultaneously.

#### Statistics

Statistical analyses were conducted using GraphPad Prism and confirmed using IBM SPSS Statistics 28.0.1 for Mac. Graphs were generated using GraphPad Prism version 9.4.1 for Mac (GraphPad Software, San Diego, CA). To examine if overall alcohol consumption was comparable in groups of males and females in this study, we first ran a three-way mixed model analysis of variance (MM-ANOVA) with one between-subjects factor and two within-subjects factors. Specifically, sex (male vs female) was the between-subject factor and DID cycle (weeks 1, 2, 3, and 4) and access length (average of three 2-hour “baseline” days vs one 4-hour “binge” day) were the within-subjects factors. There were no sex differences in alcohol intake (F _(1, 42)_ = 1.288, p = 0.2629) and no significant interactions (sex X DID cycle, F _(3, 42)_ = 1.045, p = 0.3828; sex X access length, F _(1, 42)_ = 2.132, p = 0.1517. To maximize statistical power for testing specific hypotheses, subsequent analyses were conducted separately in males and females. Data were analyzed using one-way and two-way repeated measures analysis of variance (RM-ANOVA). Bonferroni’s or Tukey’s multiple comparisons test were used for *post-hoc* analyses following significant main effects or interactions, as appropriate. Bonferroni *post-hoc* analyses were used for comparing two groups, e.g., baseline vs binge. Tukey *post-hoc* analysis were used for comparing each group with every other group (e.g., compare each DID cycle). In Experiment 1, myelin fiber density (% of area covered by MOG), density of OPCs (PDGFRɑ+ cells/mm^2^), and density of mature OLs (QKI-7+ cells/mm^2^) were analyzed using unpaired two-tailed t-tests with treatment (alcohol vs control) as the between-subject factor. Pearson’s correlations were used for simple linear regression analyses of relationships between alcohol intake in the last week and the density of myelinated axons, OPCs, and OLs. In Experiment 2, the density of pre-myelinating (QKI-7+/ASPA- cells/mm^2^), myelinating (QKI-7+/ASPA+ cells/mm^2^), and post-myelinating (QKI-7-/ASPA+ cells/mm^2^) OLs were analyzed in three anterior-to-posterior locations using two-way RM-ANOVA with treatment (alcohol vs control) as the between-subject factor with distance from bregma (AP+2.0mm vs AP+1.7mm vs AP+0.9mm) as the within-subject factor. Pearson’s correlations were used for simple linear regression analyses of relationships between alcohol intake in the last week and the density of pre-myelinating, myelinating, and post-myelinating cells. Alcohol consumption in on the last DID cycle was also used to rank alcohol drinking mice as “high” vs “low” alcohol intake groups based on a median split. Two-tailed t-tests were used to test if there were fewer QKI-7+/ASPA+ cells/mm^2^ in high drinkers compared to low drinkers. Data are presented as mean ± standard error of the mean (SEM) unless otherwise indicated. The criterion for statistical significance was *p* ≤ 0.05.

## Results

### Alcohol drinking was comparable in adolescent male and female mice

An overview of the experimental design and drinking data is shown in **Fig 1**. Beginning on PD28, the DID model was used to expose mice to two weeks of binge drinking of 20% v/v alcohol (or water for controls, **Fig. 1A**). Three days after the last drinking session, mice were perfused and brains processed for immunofluorescence experiments, followed by confocal imaging and cellular analyses. Voluntary alcohol intake was greater in the four-hour “binge” day compared to the average daily drinking during baseline days in males (**Fig. 1B**; main effect of DID cycle (F _(3, 42)_ = 23.87, p < 0.0001), access length (F _(1, 14)_ = 14.98, p = 0.0017) and cycle x length interaction (F _(3, 42)_ = 4.332, p = 0.0016)). *Post-hoc* analyses showed increased drinking on the binge day compared to the average intake on the baseline days on DID cycles 2, 3 and 4 in males (Bonferroni’s multiple comparisons test, ps = 0.0003 on week 2 and 3, and p = 0.030 on week 4). In females, alcohol consumption was greater on the binge day compared to the average baseline drinking (**Fig. 1C**), with analyses showing a main effect of access length (F _(1, 7)_ = 37.08, p = 0.0005) and cycle x length interaction (F _(3, 21)_ = 3.99, p = 0.02). *Post-hoc* analyses showed that binge day alcohol consumption was greater than the average intake on the baseline days in females on DID cycles 1, 2 and 3 (Bonferroni’s multiple comparisons test, p = 0.0007 on cycle 1, and p < 0.0001 on cycle 2 and 3).

**Figure 1.**
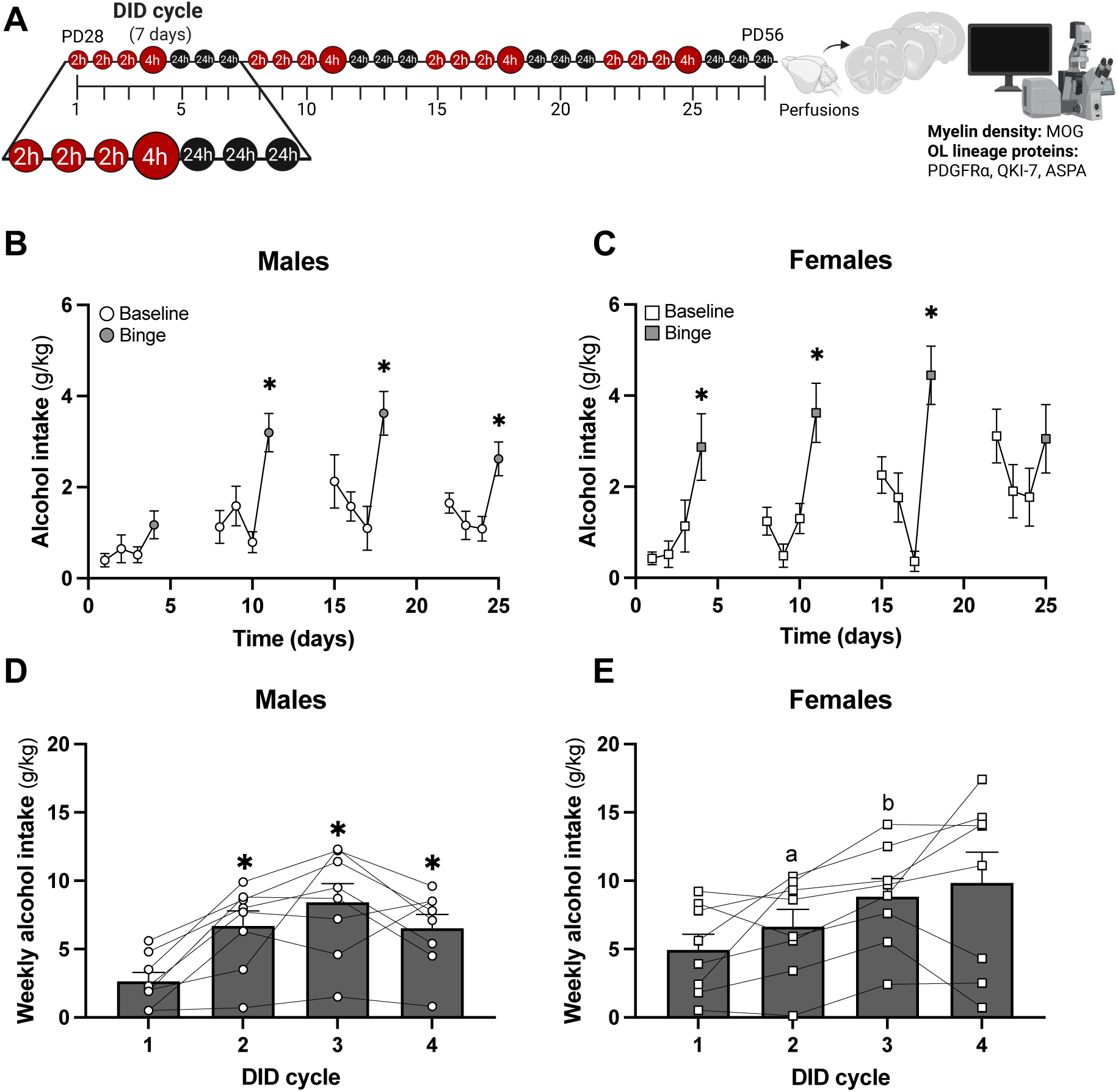
Overview of study design and alcohol intake in adolescent male and female mice. **A.** Schematic diagram of the drinking-in-the-dark (DID) alcohol binge drinking protocol. One DID cycle is three days of 2-hour “baseline” access (small red circles) to a single bottle of 20% v/v alcohol, one day of 4-hour “binge” access (big red circles) and three days with no access to alcohol (black circles). Adolescent male and female C57BL6/J mice were allowed access to alcohol for four cycles beginning on postnatal day (PD) 28. At PD56 mice were intracardially perfused and brains were processed for immunofluorescence experiments with antibodies against the indicated proteins. Sections were then imaged using confocal microscopy and analyzed. **B, C.** Daily alcohol intake (g/kg) on binge days was higher than the average of daily drinking during “baseline” days in the same DID cycle (*, all ps < 0.05, repeated measures two-way ANOVA, Bonferroni’s *post hoc*) on cycles 2, 3 and 4 in males (B) and cycles 1, 2, and 3 in females (C). **D, E.** Total weekly alcohol intake was significantly greater (*, all ps < 0.05, repeated measures one-way ANOVA, Tukey *post hoc*) in DID cycles 2, 3, and 4 compared to DID cycle 1 in males (D) and in DID cycle 3 compared to DID cycle 2 in females (E). Data are presented as mean values ± SEM; *p ≤ 0.05 = significance criterion; ns = non-significance.

Total alcohol intake in adolescent males per DID cycle showed a significant difference (F _(2.723, 19.06)_ = 11.72, p = 0.0002) and *post-hoc* analyses found a significant increase in total alcohol intake between cycle 1 and every following DID cycle (**Fig. 1D**, Tukey’s multiple comparisons test, p = 0.0093 for cycle 1 vs 2, p = 0.0031 for cycle 1 vs 3, and p = 0.0249 for cycle 1 vs 4). No significant differences were found between DID cycle 2 vs 3, cycle 2 vs 4, or cycle 3 vs 4 (all ps > 0.05). Adolescent female mice also showed a significant difference in total alcohol intake per DID cycle (F _(1.555, 10.89)_ = 4.539, p = 0.0440), and *post-hoc* analyses showed an increase in total alcohol intake only between DID cycle 2 vs 3 (**Fig. 1E**, Tukey’s multiple comparisons test, p = 0.0038). On binge days, mice surpassed alcohol consumption greater than 3 g/kg: in cycles 2 and 3 in males and in cycles 2, 3, and 4 in females. Alcohol consumption at 3 g/kg and greater is predicted to correlate with BAC over 80 mg/dl, a value that fits the criteria of alcohol binge drinking in humans (Rhodes et al., 2005b; Crabbe et al., 2009). In males, the average intake across DID cycles 3-4 is significantly greater in both baseline and binge access compared to cycles 1-2 (**Supplemental Fig. 1A**, F _(1, 7)_ = 17.28, p = 0.0005, main effect of cycle). This suggests that males increase alcohol intake across adolescent development. In contrast, females do not show a significant difference in the average intake between DID cycles 1-2 and 3-4 (**Supplemental Fig. 1B**, F _(1, 7)_ = 4.36, p = 0.07, main effect of cycle), suggesting that females consume higher amounts of alcohol from the onset of adolescence.

Water intake in control male mice was not different across the DID cycle or between baseline and binge sessions (**Supplemental Fig. 2A**, repeated measures two-way ANOVA, p > 0.05). Water intake in control female mice was significantly different on the baseline and binge days (**Supplemental Fig. 2B**, main effect of access, repeated measures two-way ANOVA, p < 0.05). No differences in the total intake per DID cycle was found in the control groups (**Supplemental Fig. 2C**, F _(1.269, 8.884)_ = 3.192, p = 0.1030 in males and **Supplemental Fig. 2D**, F _(2.214, 15.50)_ = 2.568, p = 0.1048 in females). These data support the interpretation that increased drinking on binge days is not due to longer time of access.

### Alcohol reduces myelination of anterior cingulate and corpus callosum axons in male mice, but not females

Based on our previous findings showing sex differences in prefrontal myelin loss following adolescent alcohol drinking in rats (Vargas et al., 2014; Tavares et al., 2019), the current study tested if differential sensitivity extended to male and female mice as well. We fluorescently labeled MOG—a protein that is enriched in myelin sheaths—to visualize segments of myelin ensheathing prefrontal axons extending from the front branches of the corpus callosum (forceps minor of the corpus callosum/cingulate cortex-layer VI or CC_FM_) out into the superficial layers of the cingulate cortex (Cg1; **Fig. 2A-C**). Representative images of each ROI of each treatment and sex group with MOG+ immunolabeling are shown in **Fig. 2D-E**. There was a significant reduction of myelin density in the CC_FM_ found between alcohol males compared to controls (**Fig. 2F**, unpaired two-tailed t-test, t _(14)_ = 3.975, p = 0.0014). In the anterior cingulate cortex (Cg1) there was also a significant reduction of myelin density after alcohol drinking in males (**Fig. 2F**, unpaired two-tailed t-test, t _(14)_ = 2.162, p = 0.0484). In females, there were no differences in the myelin density between the alcohol group and controls in the CC_FM_ region (**Fig. 2G**, unpaired two-tailed t-test, t _(13)_ = 1.140, p = 0.2747) nor the Cg1 region (**Fig. 2G**, unpaired two-tailed t-test, t _(13)_ = 0.5317, p = 0.6067). Given that males showed increased intake in DID cycle 4 compared to cycle 1, we tested the relationship between alcohol intake and myelin density. No significant relationships were found between the total alcohol intake of DID cycle 4 and the density of myelin sheaths in either CC_FM_ or Cg1 region of male (**Fig 2H**, simple linear regression, r^2^ = 0.06, p = 0.57 in the CC_FM_ region and r^2^ = 0.03, p = 0.66 in the Cg1 region) or female (**Fig. 2I**, r^2^ = 0.35, p = 0.16 in the CC_FM_ region and r^2^ = 0.24, p = 0.22 in the Cg1 region) mice.

**Figure 2.**
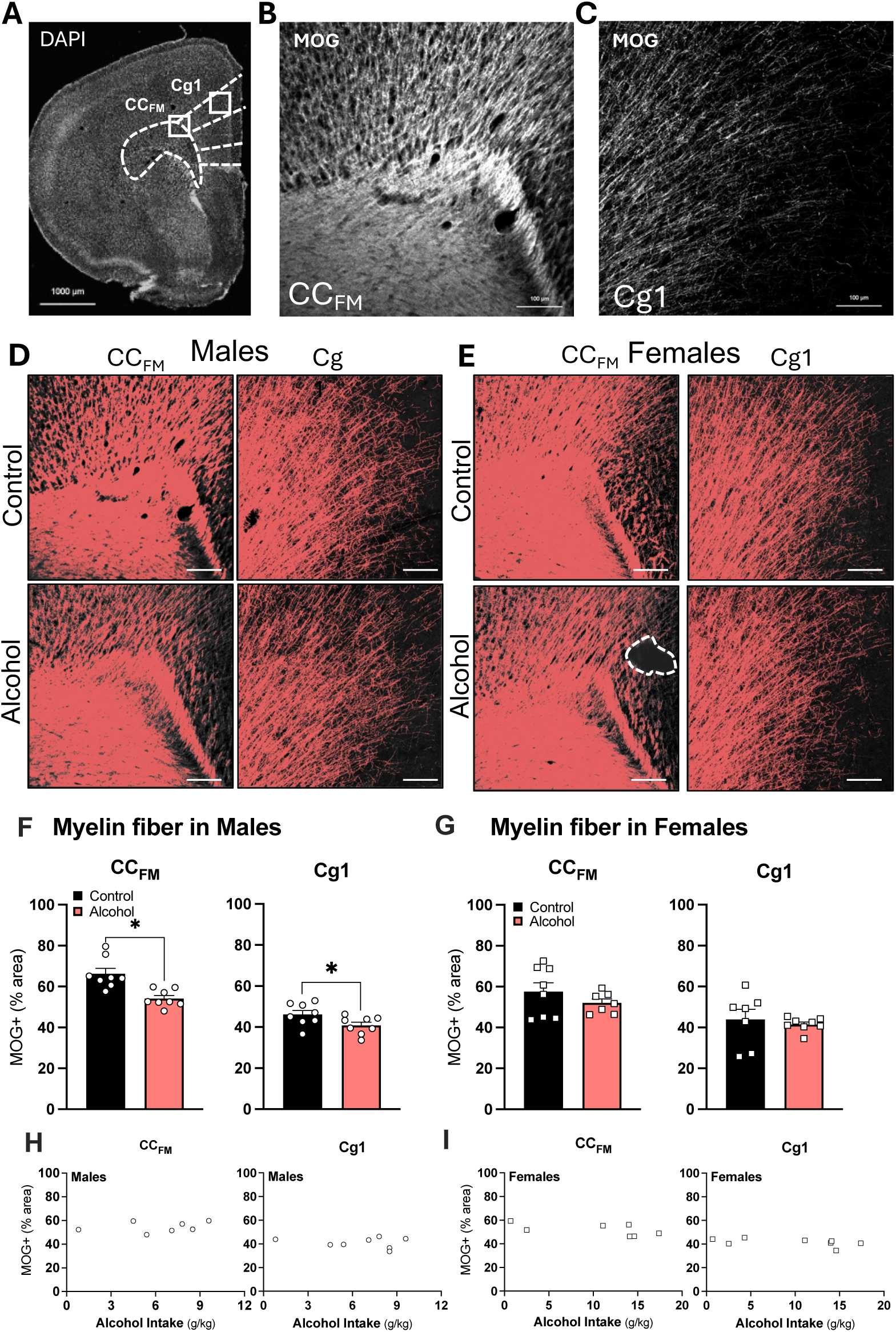
Adolescent drinking decreases the density of myelinated axons the corpus callosum and anterior cingulate in male mice. **A.** Composite image of a representative brain section with DAPI nuclear staining that was produced by stitching together frames of 10x confocal images. Sections were AP+1.7mm distance from bregma and the CC_FM_ and Cg1 sampling regions are denoted by the white boxes. Mag bar, 1000 µm**. B, C.** Representative image of the CC_FM_ (B) and Cg1 (C) acquired using the 20X objective with immunohistochemical labeling of myelin basic protein (MOG)+ myelin sheaths in white. **D, E.** Representative images showing NIS Elements thresholding of MOG+ myelin sheaths (salmon color) in the CC_FM_ and Cg1 of male (D) and female (E) control and alcohol mice. Tissue tears (dashed-line outlined area in E) were excluded from the analysis. Mag bar, 100 µm. **F, G.** Alcohol decreases myelin fiber density (% area covered by MOG) in the CC_FM_ and Cg1 regions of males (F, *, all ps ≤ 0.05, compared to controls, unpaired t-tests) but not females (G, unpaired t-tests, ps > 0.05). **H, I.** No relationship was found between the total alcohol intake in cycle 4 and myelin density in males (H) or females (I). AP, anterior-posterior position; CC_FM_, corpus callosum forceps minor; Cg1, anterior cingulate cortex; DAPI, 4’,6-diamidino-2-phenylindole; OL, oligodendrocyte, MOG, myelin oligodendrocyte glycoprotein, QKI-7, Quaking protein isoform 7. Data are presented as mean values ± SEM; *p σ; 0.05 = significance criterion; ns = non-significance.

### The pool of oligodendroglial precursor cells is not reduced by alcohol in male mice

To test the hypothesis that alcohol disrupts oligodendroglial in the early phases of cell development, thus reducing the entire OL lineage thereafter, we quantified the density of PDGFRɑ+ OPCs in the Cg1 and CC_FM_ (**Fig. 3A-E**). Contrary to our initial hypothesis, myelin deficits following alcohol drinking in males were not explained by a significant reduction in the OPC pool in these regions (CC_FM_, **Fig. 3F**, unpaired two-tailed t-test, t _(14)_ = 0.2976, p = 0.7704; Cg1, **Fig. 3F**, paired two-tailed t-test, t _(14)_ = 1.732, p = 0.1052). Likewise, alcohol did not significantly alter the OPC pool in the CC_FM_ (**Fig. 3G**, unpaired two-tailed t-test, t _(11)_ = 0.3358, p = 0.7433) or the Cg1 region (**Fig. 3G**, unpaired two-tailed t-test, t _(11)_ = 0.6082, p = 0.5554) of females. Despite a lack of significant differences in the density of OPCs in either region following alcohol, there were correlations between the level of alcohol intake in the last week of DID and OPC density in males. Using simple linear regression analyses, we detected a modest but significant negative relationship between alcohol intake and OPC density in the Cg1 region (**Fig. 3H**, r^2^ = 0.60, p = 0.03) and a modest positive significant relationship between these variables in the CC_FM_ region (**Fig. 3H**, r^2^ = 0.56, p = 0.03) and of male mice. Thus, higher intake in male mice correlated with lower OPC density in gray matter and higher OPC density in white matter, possibly indicating regional differences in the rate and timing of oligodendrogenesis changes that were captured four days after drinking ended. No significant correlations between these two variables were found in females (**Fig. 3I**, simple linear regression, r^2^ = 0.10, p = 0.53 in the CC_FM_ region and r^2^ = 0.08, p = 0.59 in the Cg1 region).

**Figure 3.**
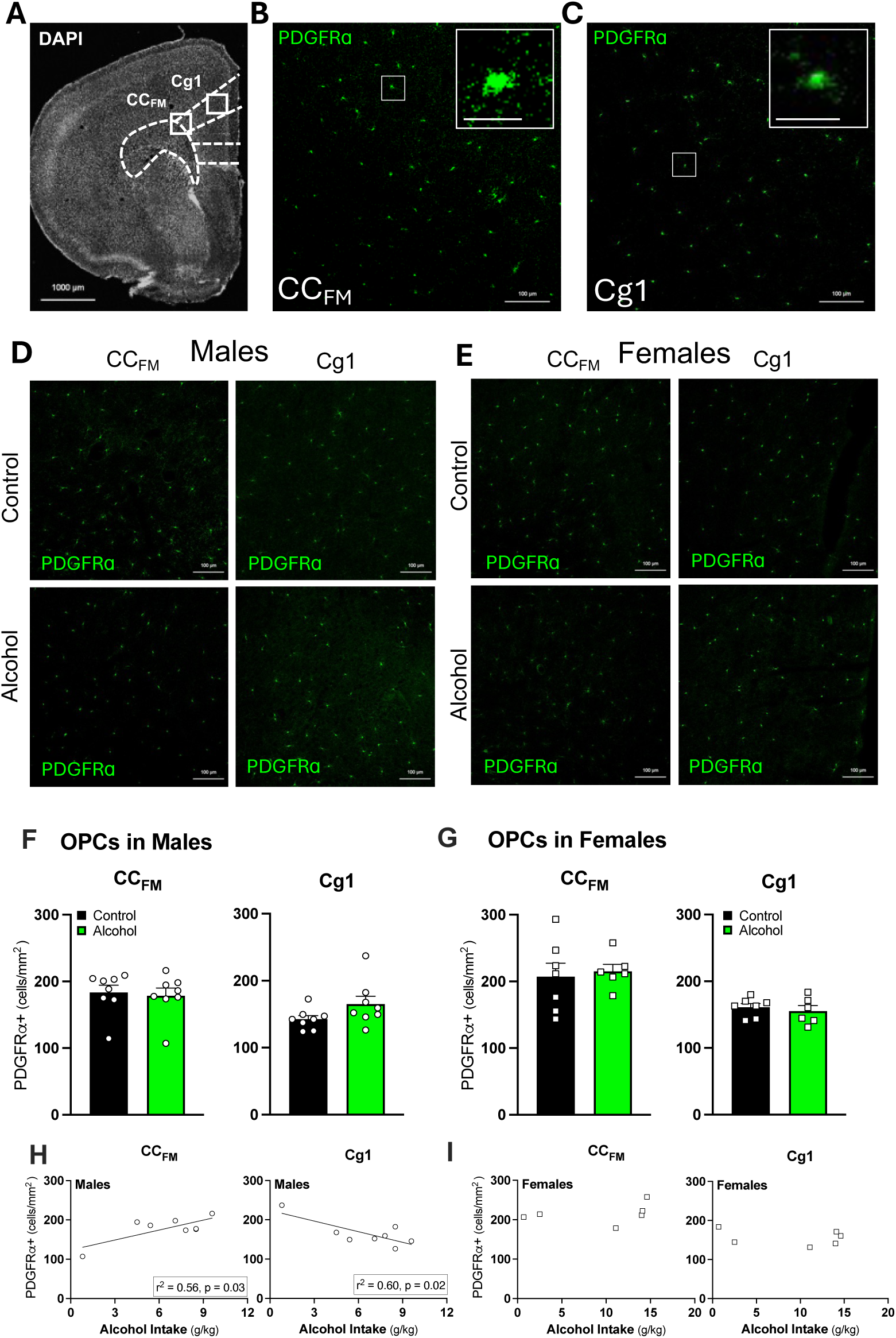
Adolescent drinking does not reduce the density of oligodendrocyte precursor cells in the corpus callosum and anterior cingulate in mice. **A.** Composite image of a representative brain section with DAPI nuclear staining, which was produced by stitching together frames of 10x confocal images. Sections were AP+1.7mm distance from bregma and the CC_FM_ and Cg1 sampling regions are denoted by the white boxes. **B, C.** Representative images of the CC_FM_ (B) and Cg1 (C) acquired using the 20X objective with immunohistochemical labeling of PDGFRɑ-expressing OPCs in green. **D, E.** Representative images of PDGFRɑ cells in the CC_FM_ and Cg1 of a control male, alcohol male, control female, and alcohol female. **F, G.** Alcohol did not affect the density of PDGFRɑ+ cells/mm^2^ in the CC_FM_ or Cg1 of males and females. **H, I.** In males, there was a significant positive relationship between the total alcohol intake in cycle 4 and OPC density in the CC_FM_ region (H, r^2^ = 0.56, *p* < 0.05) and in the Cg1 region there was a significant negative relationship (I, r^2^ = 0.60, *p* < 0.05). No significant correlations were found in females. AP, anterior-posterior position; CC_FM_, corpus callosum forceps minor; Cg1, anterior cingulate cortex; DAPI, 4’,6-diamidino-2-phenylindole; OL, oligodendrocyte, MOG, myelin oligodendrocyte glycoprotein, QKI-7, Quaking protein isoform 7. Data are presented as mean values ± SEM; *p σ; 0.05 = significance criterion; ns = non-significance. Scale bars = 1000 µm (A), 100 µm (B-E), 20 µm (inset images in B and C).

### The density of mature oligodendrocytes is increased by alcohol in the anterior cingulate

Using the well-established CC1 antibody to detect the RNA-binding QKI-7 protein as an marker for mature OLs, we tested for a decrease in OL maturation in binge drinking males (**Fig. 4A-E)**. We first confirmed the QKI-7+ cells had fully differentiated by the absence of PDGFRɑ+ signal in the cells (*data not shown*). Contrary to our prediction, mature QKI-7+ cell density was not significantly reduced by alcohol in the CC_FM_ region in males (**Fig. 4F**, unpaired two-tailed t-test, t _(14)_ = 0.8772, p = 0.3952). Instead, we found a trend of an *increase* in the number of mature oligodendroglia in the Cg1 after drinking in males (**Fig. 4F**, unpaired two-tailed t-test, t _(14)_ = 2.014, p = 0.0636), and this alcohol-induced increase was significant in females (**Fig. 4G**, unpaired two-tailed t-test, t _(11)_ = 2.929, p = 0.0137). Similar to males, the density of mature OLs was not impacted by alcohol in the CC_FM_ region (**Fig. 4G**, paired two-tailed t-test, t _(11)_ = 0.1475, p = 0.8854). The increase in QKI-7+ cell density in the Cg1 region in females and a trend of an increase in males four days after alcohol could be reflecting increases in differentiation and maturation of OLs in response to myelin loss, albeit an unsuccessful attempt at rescuing myelin deficits in males. Given the differential levels of alcohol consumption, we investigated whether total alcohol intake during the last DID cycle predicted the density of mature OLs. No significant correlations were found in males (**Fig. 4H**, simple linear regression, r^2^ = 0.16, p = 0.32 in the CC_FM_ region and r^2^ = 0.05, p = 0.60 in the Cg1 region) or in females (**Fig. 4I**, simple linear regression, r^2^ = 0.15, p = 0.44 in the CC_FM_ region and r^2^ = 0.32, p = 0.44 in the Cg1 region).

**Figure 4.**
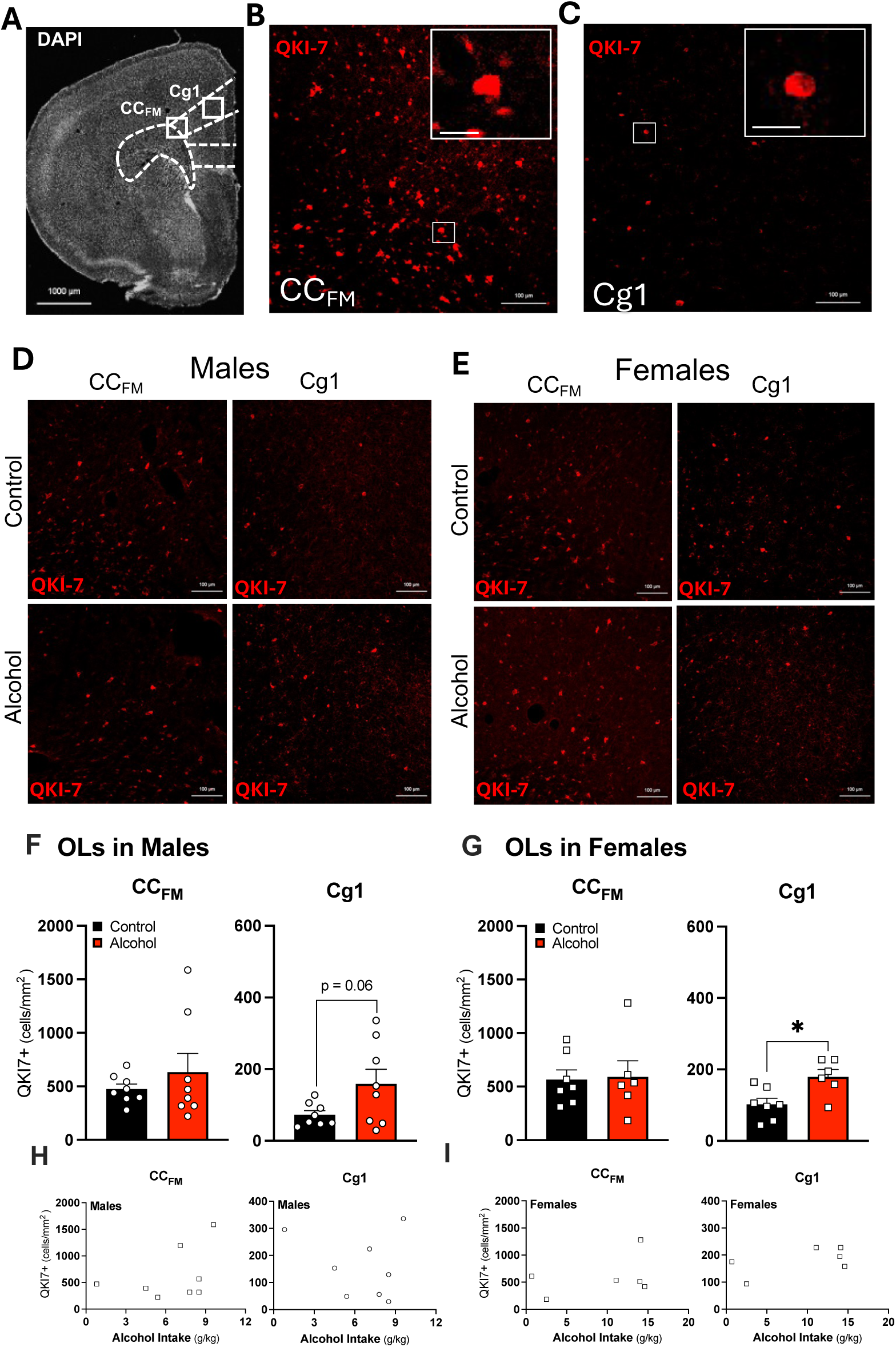
Adolescent drinking increased the density of mature oligodendroglia in the anterior cingulate in female mice. **A.** Composite image of a representative brain section with nuclear staining (DAPI) that was produced by stitching together frames of 10x confocal images. Sections were AP+1.7mm distance from bregma and the CC_FM_ and Cg1 sampling regions are denoted by the white boxes. **B, C.** Representative image of the CC_FM_ and Cg1 acquired using the 20x objective with immunohistochemical labeling of mature oligodendrocytes with the QKI-7 clone-CC1 antibody in red. **D, E.** Representative images of QKI-7 immunohistochemical labeling in the CC_FM_ and Cg1 of control and alcohol groups in males (D) and females (E). Mag bar, 100 µm. **F.** No differences were found in the density of mature oligodendroglia between alcohol drinking group and controls in males in the CC_FM_ region (unpaired t-test, p>0.05) and a trend of increase in the Cg1 region (unpaired t-test, p = 0.06). **G.** No changes in the density of differentiated oligodendrocytes were found in the CC_FM_ region in females (unpaired t-test, p>0.05); in contrast, alcohol drinking during adolescence increased the density of differentiated oligodendroglial cells in the Cg1 region in female mice (*p < 0.05, compared to controls, unpaired t-test). **H, I.** No relationship between the total alcohol intake in cycle 4 and the density of mature oligodendrocytes was found in males (H) or females (I). Data are presented as mean values ± SEM; *p ≤ 0.05 = significance criterion; ns = non-significance. Scale bars = 1000 µm (A), 100 µm (B-E), 20 µm (inset images in B and C).

### Alcohol decreased the density mature OLs expressing aspartoacylase in male mice

While the pool of QKI-7 expressing OLs appeared normal or even elevated in binge drinking males, we reasoned that there may be differences among this population that could explain hypomyelination of prefrontal axons in these animals. One possibility is that alcohol may disrupt the ability of mature OLs to myelinate. To test this possibility, we co-immunolabeled sections with both QKI-7+ and aspartoacylase (ASPA) to distinguish between mature OLs that were in a pre-myelinating stage from those that were forming myelin sheaths (**Fig. 5**). ASPA catalyzes deacetylation of N-acetyl aspartate (NAA) into free acetate, a precursor necessary for the synthesis of lipids used for myelin sheaths (Madhavarao et al., 2002). Thus, if alcohol was interfering with late cellular maturational stages in OLs they may be kept at the pre-myelinating stage, preventing the lipid synthesis necessary for the formation of myelin resulting in hypomyelination. Using ASPA as a proven specific marker for actively myelinating OLs (Pan et al., 2020), we phenotyped mature OLs into ASPA+ vs ASPA- to examine their capacity to synthesize myelin sheaths. We tested the following predictions: 1) alcohol would decrease in density of myelinating OLs (QKI-7+/ASPA+ cells) in male mice, and 2) this would be accompanied by an increase in the density of pre-myelinating OLs (QKI-7+/ASPA- cells) indicating that mature OLs were stuck at this late maturational stage of oligodendrogenesis.

**Figure 5.**
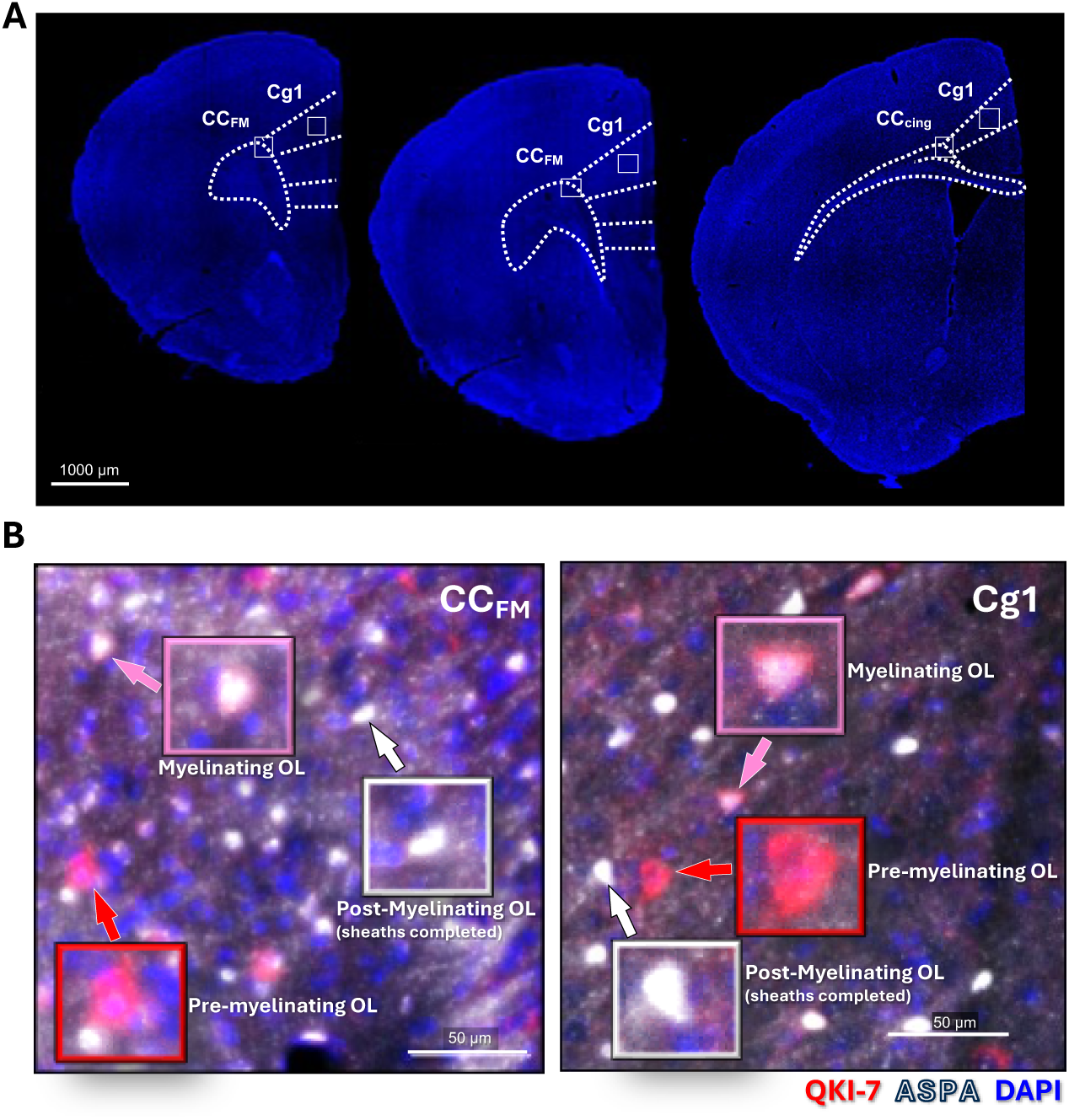
Phenotyping mature oligodendrocytes. **A.** Composite images of representative brain sections were produced by stitching together frames of 10x confocal images of a control female mouse. Sections were analyzed at AP+2.0 mm, AP+1.7 mm, and AP+0.9 mm distances from bregma, with sampled regions of the CC_FM_ and Cg1 denoted by white boxes. **B.** Representative 20x magnification images showing QKI-7 (red) and ASPA (white) immunofluorescent labeling and DAPI nucleic acid dye counterstain (blue) in the CC_FM_ and Cg1 of a control female mouse (AP+1.7mm). Three distinct OL populations were identified: pre-myelinating OLs (QKI-7+/ASPA-, red arrow), myelinating OLs (QKI-7+/ASPA+, pink arrow), and post-myelinating OLs (QKI-7-/ASPA+, white arrow). AP, anterior-posterior position; ASPA, aspartoacylase, DAPI, 4’,6-diamidino-2-phenylindole; OL, oligodendrocyte, QKI-7, Quaking protein isoform 7. Scale bars = 1000 µm (A), 50 µm (B).

As myelin deficits were observed in the medial prefrontal cortical regions (bregma AP+1.7mm), but a single myelinating OL can generate between 20 and 60 myelinating processes with intermodal lengths of about 20 mm–200 mm (Simons and Nave, 2016). Thus, we expanded our sampling area at distance range from bregma AP+2.0mm to AP+0.9mm to capture all myelinating OLs within reach of AP+1.7mm (**Fig. 5A**). There were three distinct populations of mature OLs (**Fig. 5B**): pre-myelinating OLs (QKI-7+/ASPA-, *red*), myelinating OLs (QKI-7+/ASPA+, *pink*) and a third –albeit smaller– population of QKI-7-/ASPA+ OLs we call “post-myelinating” (*white*). Representative images of AP+1.7mm for mice in the control and alcohol group are shown in **Fig. 6A**.

**Figure 6.**
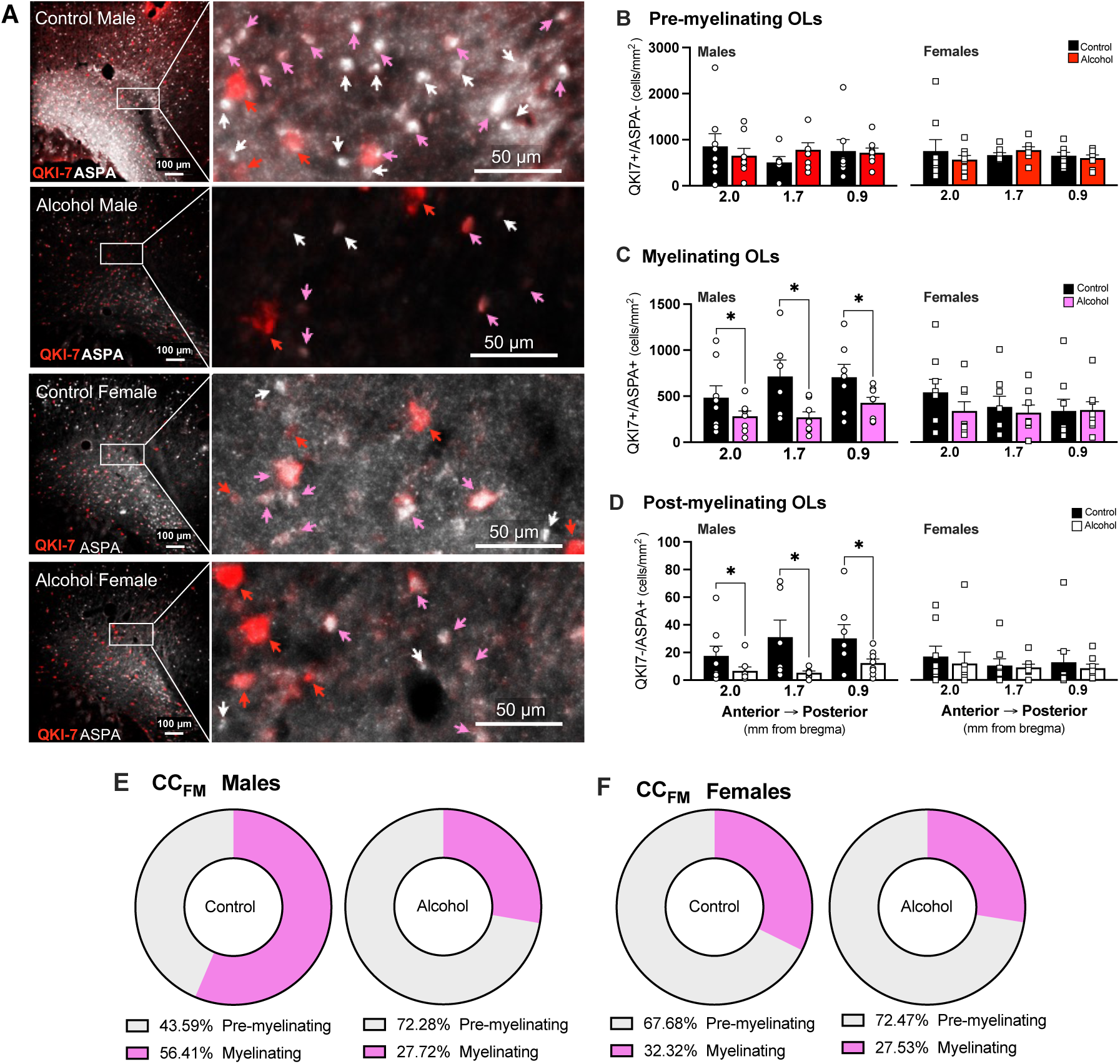
Alcohol decreases the population density of myelinating oligodendrocytes in the CC_FM_ of male mice. **A.** Representative 20x confocal images of the CC_FM_ region at AP+1.7mm from bregma in male and female mice following adolescent drinking of alcohol or water. Insets (white boxes) show higher magnification views; red arrows indicate cells expressing QKI-7 (pre-myelinating OLs), white arrows indicate cells expressing ASPA (myelinating OLs), and pink arrows indicate cells expressing both QKI-7 and ASPA (post-myelinating OLs). **B.** Pre-myelinating OL density was not affected by alcohol in either males or females (two-way ANOVAs, ns). **C, D.** Alcohol decreased the density of myelinating (C) and post-myelinating (D) OLs in males only (two-way ANOVAs, *p < 0.05, main effect of treatment in males; ns in females). **E, F.** Alcohol increased the proportion of pre-myelinating OLs in the CC_FM_ of males (E, unpaired t-test, p < 0.05), but not females (F, unpaired t-test, ns) at AP+1.7mm. Data are presented as mean values ± SEM with individual values shown in circles (males) or squares (females); *p ≤ 0.05 = significant; ns = not significant. Scale bars = 100 µm (A), 50 µm (A, close-up images).

In the CC_FM_ region of males, no significant difference in the density of pre-myelinating OLs was found between the control and alcohol groups (**Fig. 6B**, two-way MM-ANOVA, F _(1, 14)_ = 0.01331, p = 0.9098) and no significant differences were found between AP distances from bregma (**Fig. 6B**, two-way MM-ANOVA, F _(2, 25)_ = 0.7114, p = 0.4992). Similarly to males, the population density of mature pre-myelinating OLs in the CC_FM_ region in female mice showed no significant change with alcohol drinking during adolescence (**Fig. 6B**, two-way MM-ANOVA, F _(1, 14)_ = 0.1648, p = 0.6969) nor any differences between AP distances from bregma were found (**Fig. 6B**, two-way MM-ANOVA, F _(2, 28)_ = 0.3294, p = 0.7221).

The density of mature myelinating OLs in the CC_FM_ region of males was significantly lower in mice that consumed alcohol during adolescence compared to controls (**Fig. 6C**, two- way MM-ANOVA, main effect of treatment, F _(1, 14)_ = 6.174, p = 0.0262). The population of mature myelinating OLs (QKI-7+/ASPA+) in the CC_FM_ region of males was found to be significantly different between AP distances from bregma (**Fig. 6C**, two-way MM-ANOVA, F _(2, 25)_ = 5.648, p = 0.0095). Follow-up analysis indicated that the average myelinating OL density in the AP+2.0mm was significantly lower when compared to AP+0.9mm (Tukey’s *post-hoc*, p = 0.0069). No differences were found between AP+2.0mm and AP+1.7mm (Tukey’s *post-hoc*, p = 0.2298) nor between AP+1.7mm and AP+0.9mm (Tukey’s *post-hoc*, p = 0.2971). In contrast to males, we found no significant differences between alcohol and control groups in the density of mature myelinating OLs in the CC_FM_ region of female mice (**Fig. 6C**, two-way MM-ANOVA, F _(1, 14)_ = 03654. p = 0.5552). No significant differences were found in the cell density of myelinating OLs between AP distances from bregma (**Fig. 6C**, two-way MM-ANOVA, F _(2, 28)_ = 1.375, p = 0.2694) in the CC_FM_ region of females.

Alcohol consumption during adolescence significantly decreased the density of mature post-myelinating OLs in the CC_FM_ region of males (**Fig. 6D**, two-way MM-ANOVA, main effect of treatment, F _(1, 14)_ = 6.415, p = 0.0239). The density of post-myelinating OLs did not differ across the anterior-to-posterior locations sampled (**Fig. 6D**, two-way MM-ANOVA, F _(2, 25)_ = 2.489, p = 0.1033). In females, we found no differences in the population of post-myelinating OLs in the CC_FM_ between alcohol and control groups (**Fig. 6D**, two-way MM-ANOVA, F _(1, 14)_ = 0.2725, p = 0.6098) or across the anterior-to-posterior locations (**Fig. 6D**, two-way MM-ANOVA, F _(2, 28)_ = 0.5079, p = 0.6072). Adolescent alcohol drinking increased the proportion cells that remained in the pre-myelinating state in the CC_FM_ region of males (44% vs 72% in pre-myelinating OLs, **Fig. 6E**, unpaired t-test, p = 0.02), whereas the shift in females was marginal (68% vs 73% in pre-myelinating OLs, **Fig. 6F**, unpaired t-test, p > 0.05).

### Alcohol decreased the density of post-myelinating OLs in the Cg1 region in male mice

The population density of pre-myelinating, myelinating, and post-myelinating OLs was assessed in the anterior cingulate cortex (Cg1) at the same three anatomical locations as described above for the corpus callosum white matter analyses (AP+2.0mm, AP+1.7mm, and AP+0.9mm, **Fig. 5A**, outlined by a white box). Representative images of cells at AP+1.7mm for male and female mice in the control and alcohol groups are shown in **Fig. 7A**. In the Cg1 region of males, no significant differences in the average density of pre-myelinating OLs were found between alcohol and control groups (**Fig. 7B**, two-way MM-ANOVA, F _(1, 14)_ = 0.03337, p = 0.8577) or across the three AP distances from bregma (**Fig. 7B**, two-way MM-ANOVA, F _(2, 25)_ = 0.5619, p = 0.5772). In female mice, we found no significant effect of alcohol in the population density of mature pre-myelinating OLs (**Fig. 7B**, two-way MM-ANOVA, F _(1, 14)_ = 0.030, p = 0.8647). There were also no significant differences across the AP distances from bregma (**Fig. 7B**, two-way MM-ANOVA, F _(2, 27)_ = 1.078, p = 0.3544).

**Figure 7.**
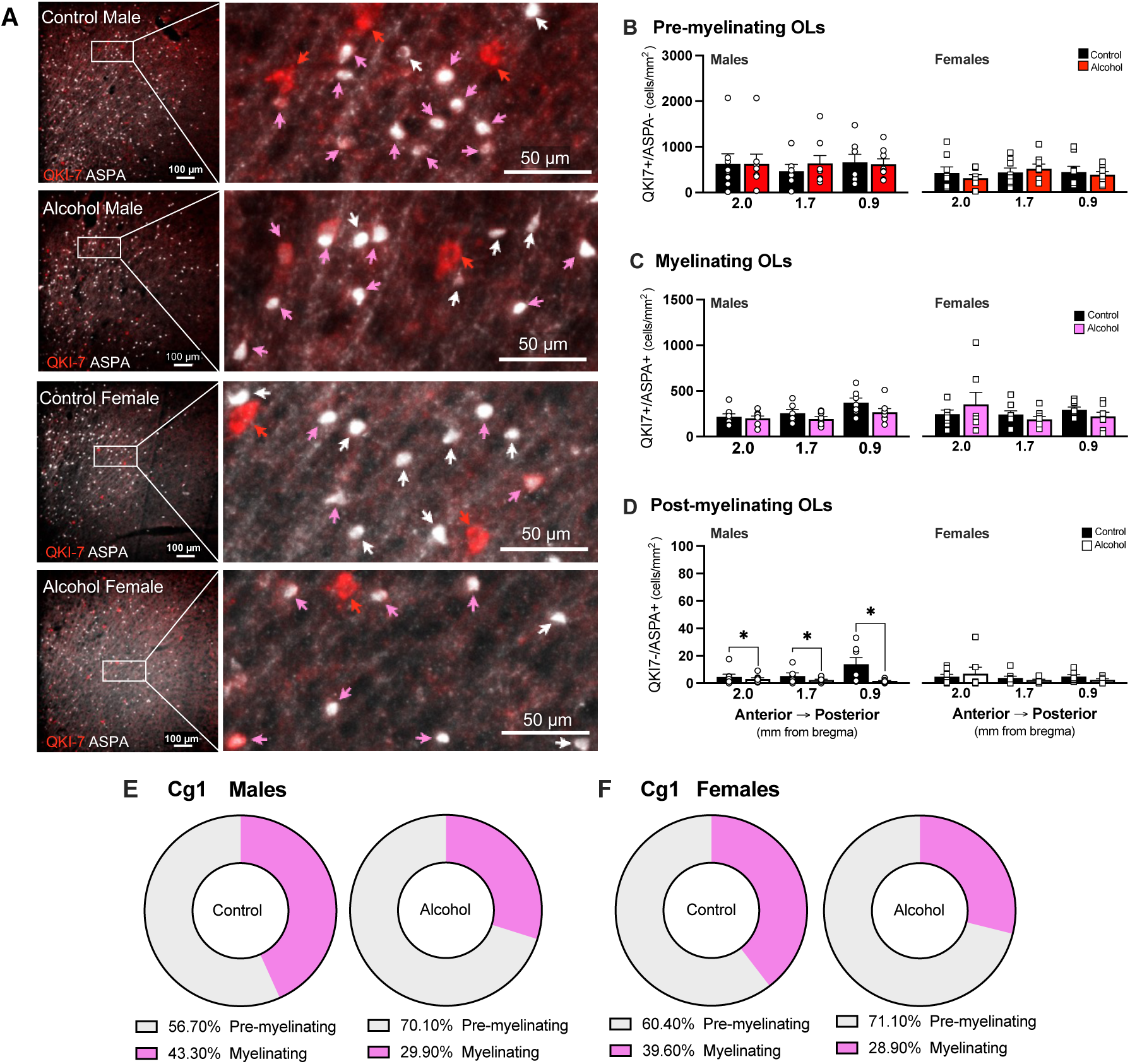
Alcohol decreases the population density of myelinating oligodendrocytes in the Cg1 region. **A.** Representative 20x confocal images of the Cg1 region at AP+1.7mm from bregma in male and female mice following adolescent drinking of alcohol or water. Zoomed in images are outlined in white boxes. Red arrows indicate QKI-7 cell, white arrows indicate ASPA cells, and pink arrows indicate both QKI-7 and ASPA cells. **B, C.** No differences found in the density of pre-myelinating and myelinating OLs in males or females (two-way ANOVAs, ns). **D.**Alcohol decreased the density of post-myelinating OLs in males, but not females (*, p < 0.05, main effect of treatment, two-way ANOVA). **E, F.** No significant changes were found in the proportion of pre-myelinating OLs in the Cg1 region in males (E, unpaired t-test, p > 0.05) or females (F, unpaired t-test, p > 0.05). Mag bar, 100 µm. Mag bars of the close-up images are 50 µm. Data are presented as mean values ± SEM with individual values shown in circles (males) or squares (females); *p σ; 0.05 = significant; ns = not significant.

No significant differences were found between alcohol and control groups in the average density of mature myelinating OLs in the Cg1 region (**Fig. 7C**, two-way MM-ANOVA, F _(1, 14)_ = 2.647, p = 0.1261); however, there was a significant main effect of AP distance from bregma (**Fig. 7C**, two-way MM-ANOVA, F _(2, 25)_ = 9.953, p = 0.0007). The average mature myelinating OL density in AP+0.9mm was significantly higher compared to AP+2.0mm (**Fig. 7C**, Tukey’s *post-hoc*, p = 0.0010) and compared to AP+1.7mm (**Fig. 7C**, Tukey’s *post-hoc*, p = 0.0049). In females, there was no effect of alcohol on the population density of mature myelinating OLs in the Cg1 region (**Fig. 7C**, two-way MM-ANOVA, F _(1, 14)_ = 0.008, p = 0.9295) and no significant differences across the three AP distances from bregma (**Fig. 7C**, two-way MM-ANOVA, F _(2, 27)_ = 1.136, p = 0.3359).

In male mice, adolescent drinking decreased the average density of post-myelinating OLs in the Cg1 region, similar to what was observed in the CC_FM_ (**Fig. 7D**, two-way MM-ANOVA, main effect of treatment, F _(1, 14)_ = 6.247, p = 0.0255). There was also a significant interaction between treatment and anatomical location (mm distance from bregma) (**Fig. 7D**, two-way MM-ANOVA, F _(2, 25)_ = 4.495, p = 0.0215), with follow-up analyses showing a significant decrease in the average density of post-myelinating OLs in the alcohol group compared to controls at AP+0.9mm distance from bregma (**Fig. 7D**, Bonferroni’s *post-hoc*, p = 0.0015) in the Cg1 region of males. No significant differences between alcohol and control groups were found at AP+2.0mm (Bonferroni’s *post-hoc*, p > 0.9999) nor AP+1.7mm (Bonferroni’s *post-hoc*, p = 0.7311). We found no significant difference in the population density of mature post-myelinating OLs in the Cg1 region in females that consumed alcohol during adolescence compared to controls (**Fig. 7D**, two-way MM-ANOVA, F _(1, 14)_ = 0.049, p = 0.8280) and no differences were found between distances from bregma (**Fig. 7D**, two-way MM-ANOVA, F _(2, 27)_ = 1.345, p = 0.2774). The proportion of the population of pre-myelinating OLs vs myelinating OLs in the Cg1 region did not significantly change with alcohol in males (**Fig. 7E**, 57% vs 70% in pre-myelinating OLs, unpaired t-test, p > 0.05) or in females (**Fig. 7F**, 60% vs 71% in pre-myelinating OLs, unpaired t-test, p > 0.05).

There were no changes in the density of mature OLs in gray or white matter regions of males or females (**Fig. 6B, 7B, and Supplemental Fig. 4A**). While there was a significant loss of myelinating OLs in the CC_FM_ region and of post-myelinating OLs in both CC_FM_ and Cg1 regions of males (**Fig. 6C, 6D, 7D, and Supplemental Figure. 4B**), there was no change in the total population of mature OLs (all QKI-7+ and ASPA+ cells, **Supplementary Fig. 4C**). This suggests that there may be a cellular compensation mechanism occurring in these regions that allow the total population of mature OLs to remain stable following alcohol.

### Alcohol intake during the last week of drinking predicts myelinating OL density in male mice

We noted a bimodal distribution of total intake consumed during DID cycle 4 in males and females (**Fig. 1D and E**). Accordingly, we used a median split to sub-divide mice into two alcohol intake groups: high drinkers (total intake > 7.4 g/kg in males and 12.6 g/kg in females; n = 4 / sex) and low drinkers (total intake < 7.4 g/kg in males and 12.6 g/kg in females; n = 4 / sex). The high intake group drank significantly more alcohol during DID cycle 4 compared to the low intake group in both males and females (**Fig. 8A**, unpaired t-tests, p = 0.02 in males and p = 0.005 in females). We next assessed whether there were significant differences in OL density between high and low drinking groups. High drinking male mice had a lower density of myelinating OLs in the Cg1 region at the distance from bregma AP+1.7mm compared to low drinking male mice (**Fig. 8B**, unpaired t-test, p = 0.0011). In contrast, the density of myelinating OLs in the Cg1 region was similar in the low and high drinking groups in female mice (**Fig. 8B**, unpaired t-test, p = 0.5722). There was also a significant negative relationship between the total alcohol intake in DID cycle 4 and the density of myelinating OLs in the Cg1 region of male mice (**Fig. 8C**, simple linear regression, r^2^ = 0.81, p = 0.0025). There was no correlation between total alcohol consumed in DID cycle 4 and the density of myelinating OLs in the Cg1 region in female mice (**Fig. 8C**, bottom scatter plot, simple linear regression, r^2^ = 0.08, p = 0.7819). When both high and low alcohol intake groups were compared to controls, the density of myelinating OLs in the Cg1 region of males was significantly decreased in the high intake group (Dunnett’s *post-hoc* following a significant one-way ANOVA, F _(2, 11)_ = 2.984, p = 0.04, *data not shown*). No differences were found in the density of myelinating OLs between the three groups in females (one-way ANOVA, F _(2, 13)_ = 0.1471, p = 0.5482, *data not shown*). These data suggest myelinating Cg1 myelinating OLs are sensitive to increasing alcohol levels in males only. This may reflect sex differences in the population of myelinating OLs during adolescent development.

**Figure 8.**
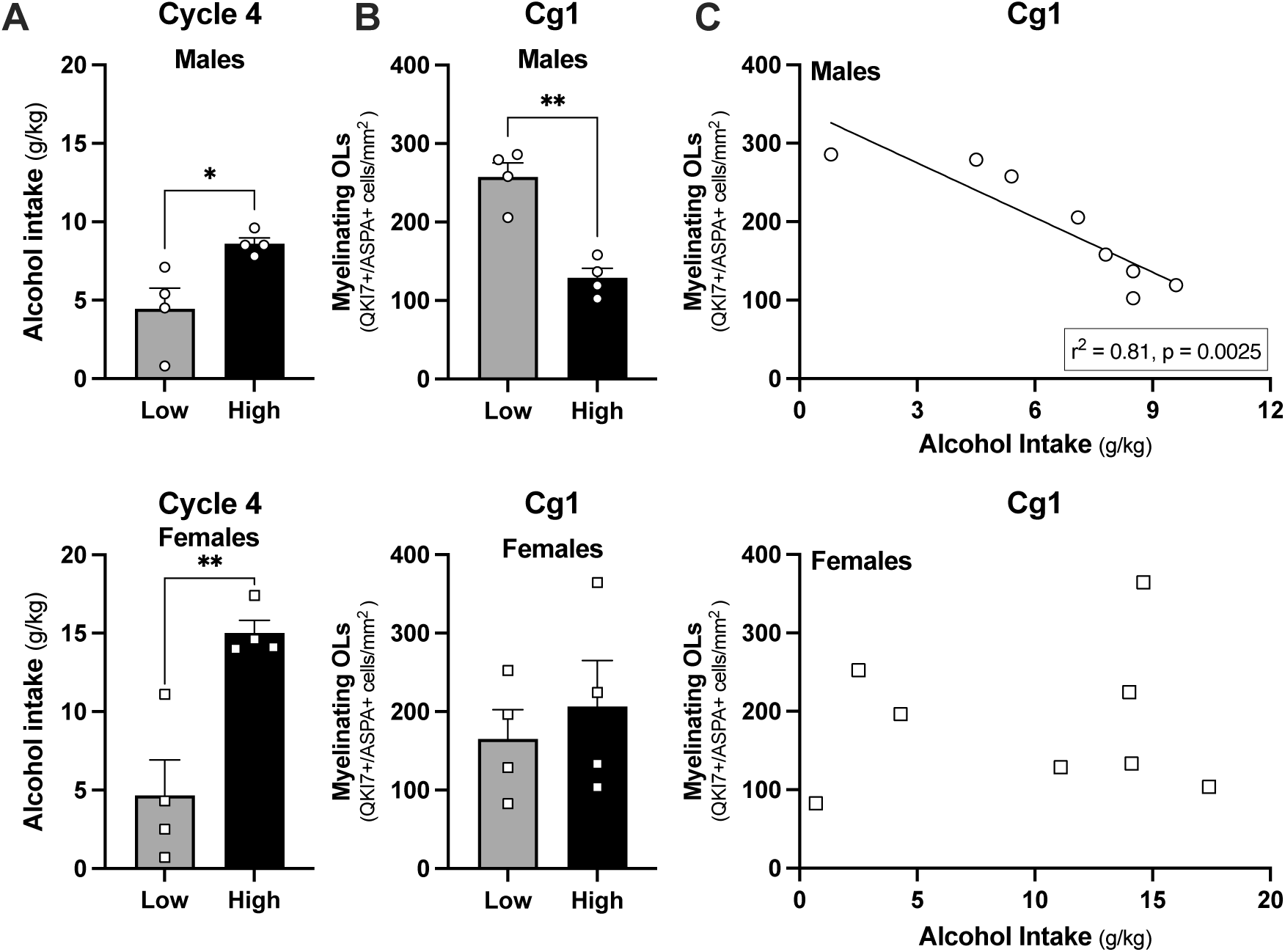
High alcohol intake predicts low myelinating OL density in the Cg1 region in males. **A.** Mice were categorized as “high” or “low” drinking groups based on a median split of the total amount of alcohol consumed during the last week of drinking (DID cycle 4, Fig 1D and E). High drinking male mice consumed twice as much alcohol as low drinking males (top bar graph, *, p < 0.05, unpaired t-test) and high drinking female mice consumed three times as much alcohol as low drinking females (bottom bar graph, **, p < 0.01, unpaired t-tests). **B.** Myelinating OL density in the Cg1 was lower in high-drinking males compared to low-drinking males in the Cg1 region at (top bar graphs, **p < 0.01, unpaired t-test). OL density was similar in high and low drinking groups of females (bottom bar graphs, p > 0.05, unpaired t-test). **C.** There was a tight negative correlation between the total amount of alcohol consumed and myelinating OL density in the Cg1 in males (top scatter plots, (r^2^ = 0.81, p < 0.01) but not females (bottom scatter plots, r^2^ = 0.08, ns). Sections used for analyses were AP+1.7mm distance from bregma. Data are presented as mean values ± SEM with individual values shown in circles (males) or squares (females); *p ≤ 0.05 = significant; ns = not significant.

**Figure 9.**
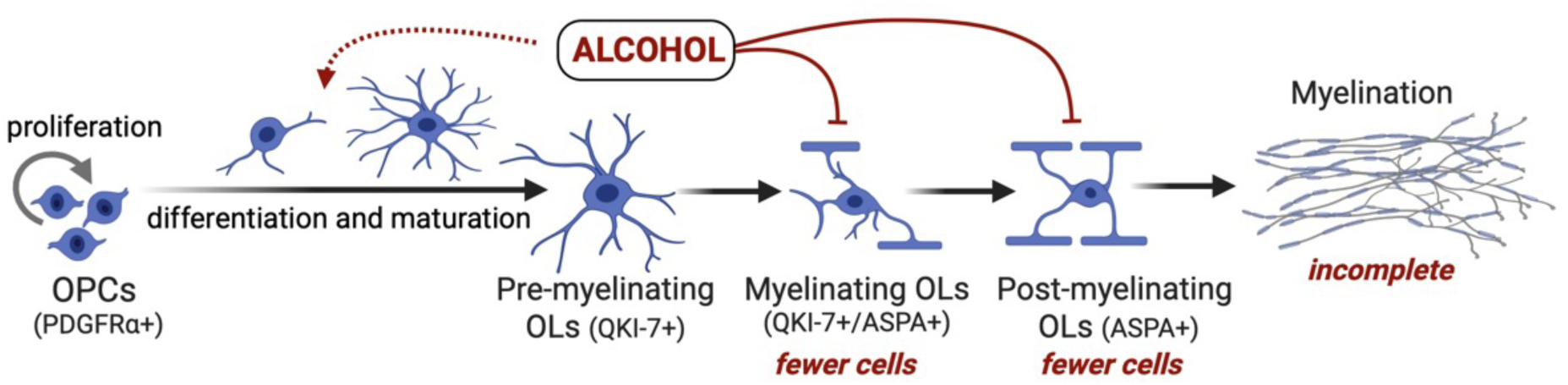
Working model illustrating how binge drinking impairs myelination of prefrontal axons during adolescent development. We propose that alcohol targets the OL lineage in the late phase of cellular development, resulting in a low number of cells expressing the ASPA enzyme that is necessary for lipid synthesis during myelination. Even if the loss of ASPA+ OLs leads to a compensatory increase in OPC differentiation, this response is insufficient to rescue the myelinating OL population in males, resulting in myelin deficits. Arrows indicate the hypothesized direct (solid lines) and indirect (dashed lines) effects of alcohol on oligodendroglia. OPC, oligodendroglia precursor cell; OL, oligodendroglial cell; PDGFR⍺, platelet-derived growth factor receptor alpha; QKI-7, quaking protein isoform-7; ASPA, aspartoacylase.

## Discussion

The present study showed that adolescent drinking disrupts myelination of axons in the anterior cingulate cortex and adjacent white matter of the corpus callosum in male mice. Our results indicate that a significant loss in mature OLs expressing ASPA may have caused hypomyelination of axons in males given the importance of this enzyme for lipid biosynthesis during myelin sheath formation (Hershfield et al., 2006; Mattan et al., 2010; Francis et al., 2016). Most notably while the magnitude of cell loss was even more pronounced in higher drinking males, females appeared resistant to the negative effects of alcohol regardless of how much they drank. Binge drinking females had a higher density of mature OLs (QKI-7+ cells) in the Cg1, and there was a similar trend in male mice. The subtle changes in OPCs and mature OL populations may signify an upregulation in oligodendrogenesis that could have replenished the ASPA+ OL pool in females, protecting them against the hypomyelinating effects of alcohol.

Nevertheless, this was clearly insufficient to fill the ASPA+ OLs pool and rescue myelin loss in alcohol males. Our results replicate previous reports of myelin deficits with alcohol in rodents and humans (Jacobus et al., 2009; Vargas et al., 2014; Papp-Peka et al., 2016; Wolstenholme et al., 2017; Rice et al., 2019; Tavares et al., 2019) and fill a significant knowledge gap by providing evidence that alcohol impacts oligodendroglial lineage cells at a later maturational stage of cellular development. By identifying ASPA as a direct or indirect target of alcohol, we highlight the need for further investigation of this enzyme, which is a promising new target for therapeutic intervention in alcohol use disorder and demyelinating diseases.

### Alcohol induces myelin sheath density loss in male mice

We previously reported that the anterior branches of the corpus callosum (CC_FM_) which project to the Cg1 region undergo substantial increases in myelin density during adolescent development, which significantly speeds up the conduction velocity in these axons (McDougall et al., 2018). These myelinated axons are vulnerable to alcohol consumption during adolescence, as alcohol reduced the density of myelinated fibers in adolescent male, but not in female, rats (Vargas et al., 2014; Tavares et al., 2019). The data from the study herein recapitulated these findings in adolescent male mice. This is consistent with evidence that alcohol is a demyelinating agent in the CNS. Previous studies measuring the gene expression of myelin-associated glycoprotein *(Mag),* myelin basic protein (*Mbp),* myelin-associated oligodendrocytic basic protein *(Mobp),* and proteolipid protein 1 *(Plp)* showed reductions in the prefrontal cortex after a bolus high dose of alcohol via gavage in adolescent mice (Wolstenholme et al., 2017). A similar high dose administration of alcohol during adolescence decreased myelin density in the prefrontal cortex, primarily in the axons of parvalbumin-negative neurons (Rice et al., 2019). These deficits in myelination during adolescence have been associated with impairments in working memory and social interaction in young adult mice (Makinodan et al., 2009).

Myelin oligodendrocyte glycoprotein (MOG) is a 28 kDa protein located in the outermost layer of the myelin sheath exclusively in the CNS and is a relatively very minor (0.05%) component of myelin (Johns and Bernard, 1999). MOG may play a role as an adhesion protein supporting myelin compaction (Clements et al., 2003). The current study shows evidence that four weeks of voluntary alcohol intake sufficiently perturbs MOG+ myelin density in male mice. Both the density of myelin sheaths and density of OLs expressing *Mog* mRNA is decreased throughout the CC and the prefrontal cortex in male mice following chronic social defeat stress (Lehmann et al., 2017). Similarly, a bolus dose of alcohol (3g/kg, *i.p*.) in female rats or chronic exposure to alcohol through continuous home-cage access for 5 months in female mice reduces *Mog* mRNA and MOG protein levels in the prefrontal cortex (Alfonso-Loeches et al., 2012; Pascual et al., 2014). Possibly longer exposure to alcohol may be necessary to induce similar deficits in females as we see in males.

### ASPA-expressing OLs are sensitive to adolescent drinking in male mice

ASPA generates the free acetate needed for lipid synthesis in myelin formation through the hydrolyzation of N-acetylaspartate (NAA) released by neurons (Madhavarao et al., 2002; Hershfield et al., 2006; Francis et al., 2012, 2016; Grønbæk-Thygesen and Hartmann-Petersen, 2024). ASPA is highly expressed in OLs, located predominantly in the soma in both the nucleus and cytoplasm, and with 92-98% co-localization with QKI-7+ mature OLs (Baslow et al., 1999; Madhavarao et al., 2004; Hershfield et al., 2006). Expression of ASPA also follows the developmental trajectory of myelination in the CNS, supporting the role of ASPA in myelin synthesis (Kirmani et al., 2003). Deficiency of ASPA enzyme, a characteristic of Canavan disease, disrupts the production of myelin-associated lipids which leads to vacuolation and myelin deficiency (Takeda et al., 2024).

High alcohol exposure has been reported to reduce brain levels of ASPA’s substrate NAA. Administration of 3 g/kg of alcohol for 4 days via *i.p.* injections decreased brain NAA levels in adolescent Swiss-Webster male mice (Baslow et al., 2000). Similarly, in a recent study on adult patients with AUD seeking treatment, NAA levels within the frontal gray and white matter were significantly lower in the high-risk to relapse group compared to light- and non-drinking controls (May et al., 2025). This may reflect a reduction in NAA synthesis in neurons after alcohol. If similar effects happen with alcohol drinking in adolescent male mice, the combination of a limited supply of the NAA substrate from neurons and the lower number of ASPA-expressing OLs to catabolize the deacetylation of NAA could conceivably cause a major deficiency in available free acetate, further exacerbating myelin loss.

It is unclear whether alcohol drinking prevented mature OLs from starting to express ASPA. Alcohol may be holding mature cells in a pre-myelinating state or may disrupt ASPA expression in actively myelinating OLs preventing them from continuing to form myelin sheaths. While the fate of these “lost” ASPA cells is unknown, the consequences of alcohol could be significant and long-lasting. Considering the possibility that the NAA substrate supplied by axons may already be lowered by alcohol as other studies have shown (Baslow et al., 2000; May et al., 2025; Sommer and Canals, 2025), and if there are also not enough mature OLs expressing ASPA to catalyze deacetylation of the NAA that is available to synthesize myelin sheaths, then drinking during this time could have lasting effects on prefrontal circuits that impact functions in adulthood.

### Differential correlations between alcohol and PDGFR+ OL progenitor cells in white and gray matter

In the present study, there were no group differences detected in OPC density in the CC_FM_ and Cg1, despite decreased density of myelinated axons and myelinating OLs in alcohol males compared to controls. This was somewhat surprising because acute cellular injury can trigger apoptotic cell death OPCs in 24 hours (Hill et al., 2017; Chapman et al., 2024) and myelin damage and OL loss accelerates the differentiation of OPC into OLs (Hill et al., 2014; Baxi et al., 2017; Chapman et al., 2023). These two events would be expected to reduce the number of OPCs, but there is a dynamic interplay between cellular division and differentiation that serves to stabilize the OPC pool. In response to an acute demyelinating event and OPC cell death, OPCs divide and a portion of daughter cells differentiate into OLs within a few days (Hill et al., 2014; Baxi et al., 2017). As brains were collected three days after the last alcohol binge day in our study, it is possible that there was enough time for the OPC pool to replenish itself through cell division after alcohol exposure ended.

Delving deeper into the OPC population, we found region-specific relationships between OPC density and the total alcohol intake on the last DID cycle. There was a modest negative correlation in gray matter, with higher drinking levels being associated with lower OPC density in the Cg1. Conversely, there was a modest positive correlation in white matter, with higher drinking levels being associated with higher OPC density in the CC_FM_. Differential dynamics of oligodendrogenesis may explain this, as OPCs proliferate and differentiate faster into OLs in white matter regions compared to gray matter following myelin injury (Baxi et al., 2017). Thus, OPC density could be elevated in the CC_FM_ of high drinking mice because proliferation was initiated earlier in the corpus callosum OPC population, as has been observed after social chronic stress (Poggi et al., 2022). In the Cg1, OPC populations may be lower due to increased differentiation in response to demyelination (Hill et al., 2014). A recently published study showed that primary culture of OLs from mice cortices have distinct gene expression changes in high (30mM, 138.3 mg/dl BAC) compared to a moderate (10mM, 46.1 mg/dl BAC) concentration of alcohol (Bazzi et al., 2025). At the moderate concentration, alcohol increased genes associated with cell cycle progression through G1 and mitosis (increase in Cyclin B and D) and decreased genes associated with progression though S and G2 (decrease in Cyclin A and E), suggesting disrupted cell division at specific steps. On the other hand, all cyclin genes were downregulated at the high concentration, suggesting a decrease in proliferation at that dose (Bazzi et al., 2025).

A modest increase in OPC density with greater alcohol intake in the corpus callosum may be due to several possible events: 1) an increase in proliferation with a failure to initiate differentiation, 2) a failure in OPC migration from white matter to the adjacent gray matter region, or 3) a white vs gray matter difference in the rate of OPC proliferation and differentiation. In support of these possible explanations, other studies have shown that OPC proliferation and differentiation into OLs occur at a greater rate in white matter compared to gray matter (Dimou et al., 2008; Rivers et al., 2008; Viganò et al., 2013; Young et al., 2013) and both proliferation and differentiation are increased following demyelination, at a faster rate in the corpus callosum compared to the cingulate cortex (Hill et al., 2014; Baxi et al., 2017). During a demyelinating insult such as cuprizone, OLs regenerate by OPC proliferation and migration during active demyelination (Mason et al., 2000). Alcohol consumption can specifically inhibit OL differentiation without changing the OPC population density (Guo et al., 2021). Furthermore, the OPCs that migrated the longest distance in response to injury were the subpopulation of progenitor cells that did not differentiate into OLs (Chapman et al., 2023). If alcohol disrupts OPC migration, this may have contributed to the region-specific changes in OPC density we observed in the present study. Therefore, these events may, in combination, compound and result in the modest increase in OPC density of the CC_FM_ region in males and a modest decrease in the Cg1 region. Our findings provide additional support for the notion that OPCs respond differently to demyelination and injury in gray versus white matter.

### Evidence for elevations in the density of QKI-7 expressing OLs after alcohol drinking

During OL maturation, these cells start producing specific proteins that are necessary for the production and maintenance of myelin sheaths (Huang et al., 2023). Both pre-myelinating and myelinating OLs express Quaking Protein-Isoform 7 (QKI-7, labeled by the CC1 antibody), a protein that binds and stabilizes the mRNA of myelin structural proteins including myelin basic protein (Bin et al., 2016). The average density of pre-myelinating OLs was not affected by alcohol in adolescent mice; however, there is a hint of change with alcohol in males. There was a trend of an increase in the density of QKI-7 OLs in the cingulate cortex. This pooling of mature OLs may indicate 1) an inability to express ASPA and generate myelin sheaths following alcohol, or 2) an enhanced OPC differentiation. Recent findings in the nucleus accumbens indicate increases in differentiation (measured as an increased number of CC1+ QKI-7 mature OLs) at 6 weeks of alcohol consumption in adult mice (Liran et al., 2025) and our findings showing a negative correlation between alcohol intake and OPC density may suggest a possible upregulation of OPC differentiation in the Cg1 of males. In mice treated with cuprizone to induce demyelination, the density of CC1+ QKI-7 OLs showed long-term (6 weeks post-treatment) increase in the corpus callosum but decrease in the cingulate cortex (Baxi et al., 2017), suggesting timing-dependent differences between the white matter and gray matter regions in the cellular response to a demyelinating event.

### Alcohol consumption was comparable between adolescent male and female mice

The drinking-in-the-dark (DID) protocol was used for alcohol administration because it reliably elicits binge-like drinking behavior and achieve biologically relevant blood alcohol concentrations (Crabbe et al., 2009; Barkley-Levenson and Crabbe, 2012; Thiele and Navarro, 2014; Thiele et al., 2014). This voluntary, limited access to alcohol allowed the mice to reach alcohol intake over 3 g/kg for both sexes. This alcohol consumption has been shown to correlate with BAC over 80 mg/dl, a value that fits the criteria of binge drinking in humans (Rhodes et al., 2005b; Crabbe et al., 2009; Thiele and Navarro, 2014; Wilcox et al., 2014). We also found higher alcohol consumption on bassline and binge days in the later DID cycles in male mice, similar to the escalated drinking patterns that have been previously reported with the DID model (Wilcox et al., 2014). Female mice showed comparable drinking to male mice, consistent with previous studies that showed similar levels of alcohol consumption between adolescent male and female mice and rats (Schramm-Sapyta et al., 2014; Tavares et al., 2019; Silva-Gotay et al., 2021; Edwards et al., 2025). Others have shown that female rodents drink more alcohol than males during adolescence (Walker et al., 2008; Strong et al., 2010) and/or in adulthood (Rhodes et al., 2005b; Walker et al., 2008; Strong et al., 2010; Flores-Bonilla et al., 2021). One study found that alcohol intake from PD30 to PD51 was comparable between male and female rats; however, a shift in increase of alcohol drinking was found in female rats compared to males from PD52 onwards (Lancaster et al., 1996). Sex differences in alcohol drinking in adulthood is driven primarily the magnitude of front-loading these animals exhibit (Flores-Bonilla et al., 2021). When adolescent rodents first start drinking, both males and females consume more alcohol than adults (Bell et al., 2006; Walker et al., 2008; Strong et al., 2010; Schramm-Sapyta et al., 2014; Lee et al., 2017). A history of adolescent drinking can lead to higher alcohol drinking later in adulthood (Gilpin et al., 2012; Pandey et al., 2015; Younis et al., 2019), and greater effects have been reported in females (Strong et al., 2010). High alcohol drinking (HAD) rat strains also show higher intake during adolescence compared to adulthood, particularly adolescent males show the highest drinking–while adolescent females show the lowest drinking– of all four age and sex groups tested (Dhaher et al., 2012). There were strain-dependent effects on alcohol drinking modulated by sex in adults, as adult HAD-1 rats did not show sex differences while adult HAD-2 female rats showed greater intake than males (Dhaher et al., 2012). Similarly, adult male alcohol-preferring rats consumed more alcohol compared to adult females (Bell et al., 2006). These differences compared to outbred strains of rats and mice may be a result of selectively breeding (for 75+ generations) for higher intake and introducing a history of excessive alcohol drinking for multiple generations that leads to blunted neuronal activity associated with decision-making related to alcohol drinking behavior (Linsenbardt et al., 2019).

### Female mice are resilient to alcohol-induced myelin loss

We did not detect measurable changes in myelinated fiber density after alcohol drinking in female mice despite exhibiting similar levels of alcohol intake using the DID alcohol binge drinking model. This is consistent with our previous report in adolescent female rats (Tavares et al., 2019). Despite studies showing reduced gene expression and protein levels of myelin-associated genes in female rodents with high doses of alcohol (Alfonso-Loeches et al., 2012; Pascual et al., 2014), these changes may be due to the methods used to expose animals to alcohol. We have found that in female rats, alcohol reduces the length of the nodes of Ranvier located between the contactin-associated protein (Caspr) pairs (Tavares et al., 2019). This has implications for action potential conduction velocity and amplitude, as a reduced nodal length would lead to a decrease in sodium channels available at the nodes (Babbs and Shi, 2013; Arancibia-Cárcamo et al., 2017; Scurfield and Latimer, 2018).

Loss of myelin density in the CC_FM_ and Cg1 regions of adolescent males but not females may reflect differences in the pubertal timing of myelination between sexes. Axons that have partial or complete myelin sheaths at the time when demyelination occurs have faster remyelination and are more selectively targeted for remyelination by OLs compared to isolated myelin sheaths (Chapman et al., 2023). Adult males have a higher density of myelin sheaths and OLs in the corpus callosum compared to females of the same age (Cerghet et al., 2006). However, this is mediated by gonadal hormones, as castrated males show lower myelin sheath density in the corpus callosum and fimbria at the anterior hippocampus compared to intact males (Cerghet et al., 2006). Since pubertal maturation occurs at an earlier age in females compared to males (Tavares et al., 2019), *de novo* myelin sheaths may be added before the start of alcohol intake in females but not males. By PD28, females may have a greater percent of axons that are partially or completely myelinated while males may have a higher percentage of isolated myelin sheaths. Since brains were collected four days post-alcohol (PD56), this allows enough time for partial and completed myelin sheaths to remyelinate while isolated myelin sheaths take approximately eight days or more for remyelination to occur (Chapman et al., 2023). Ongoing studies are dissecting the sex differences in the rate of myelination during early adolescent development.

No change in the population of OPCs, pre-myelinating, myelinating, and post-myelinating were found in the cingulate cortex or corpus callosum of female mice in this study. However, alcohol increased the density of mature (QKI-7+ OLs) in the cingulate cortex of female mice, indicating a potential mechanism that ultimately results in the preservation of myelin sheaths. Adult females rodents have a lower myelin and OL density and a higher turnover (increased proliferation and cell death) of OLs in the corpus callosum, fornix, and spinal cord compared to males that is mediated by gonadal hormones (Cerghet et al., 2006). In support of this, administration of 17-β estradiol (alone or combined with progesterone) in male mice partially blunts the effects of cuprizone on OL density loss and demyelination in the corpus callosum (Acs et al., 2009; Taylor et al., 2010). This hormone-mediated ability for greater rate of OL replacement suggests that females may have a mechanism for faster renewal of myelin in response to insults like alcohol. This is the case for 12-month-old female rats following a demyelinating lesion induced by ethidium bromide injection (Li et al., 2006). However, they also found no differences in the remyelination rate in young adult (2-month-old) between male and female rats (Li et al., 2006). Whether sex-specific differences in alcohol-induced myelin loss is due to 1) timing of exposure relative to the surge of gonadal hormones, 2) sex differences in OL renewal and/or remyelination rate, or 3) hormonal neuroprotection conferring resiliency to OLs in females is still a subject of active research.

### Conclusions

We have demonstrated that myelinating (ASPA+) OLs are particularly vulnerable to alcohol in adolescent male mice, shifting the dynamics of differentiation and maturation of the oligodendrocyte lineage, resulting in loss of myelin sheaths. These results contribute to the growing body of evidence that alcohol disrupts the maturation of frontotemporal circuits, leading to delayed processing and both functional and behavioral consequences, increasing the risk of developing AUD later in life.

## Acknowledgements

We kindly thank Rithika Senthilkumar and Faye Reagan for their assistance with these studies, and the university animal husbandry and veterinary staff for their dedicated support in maintaining the health and welfare of our mice. We performed confocal imaging and analyses and imaging in the UMass IALS Light Microscopy Facility and Nikon Center of Excellence, and we thank the Director, Dr. James Chambers, for his support and guidance. The work was funded with support from the National Institutes of Health R01AA024774 (HNR) and NIH R21AA031376 (HNR), the National Science Foundation Graduate Fellowship Research Program 1938059 (AFB), and the UMass Spaulding-Smith Fellowship Program (AFB). Graphs were created with GraphPad PRISM and figures were created with BioRender.com.

---

**Supplemental Figure 1.**
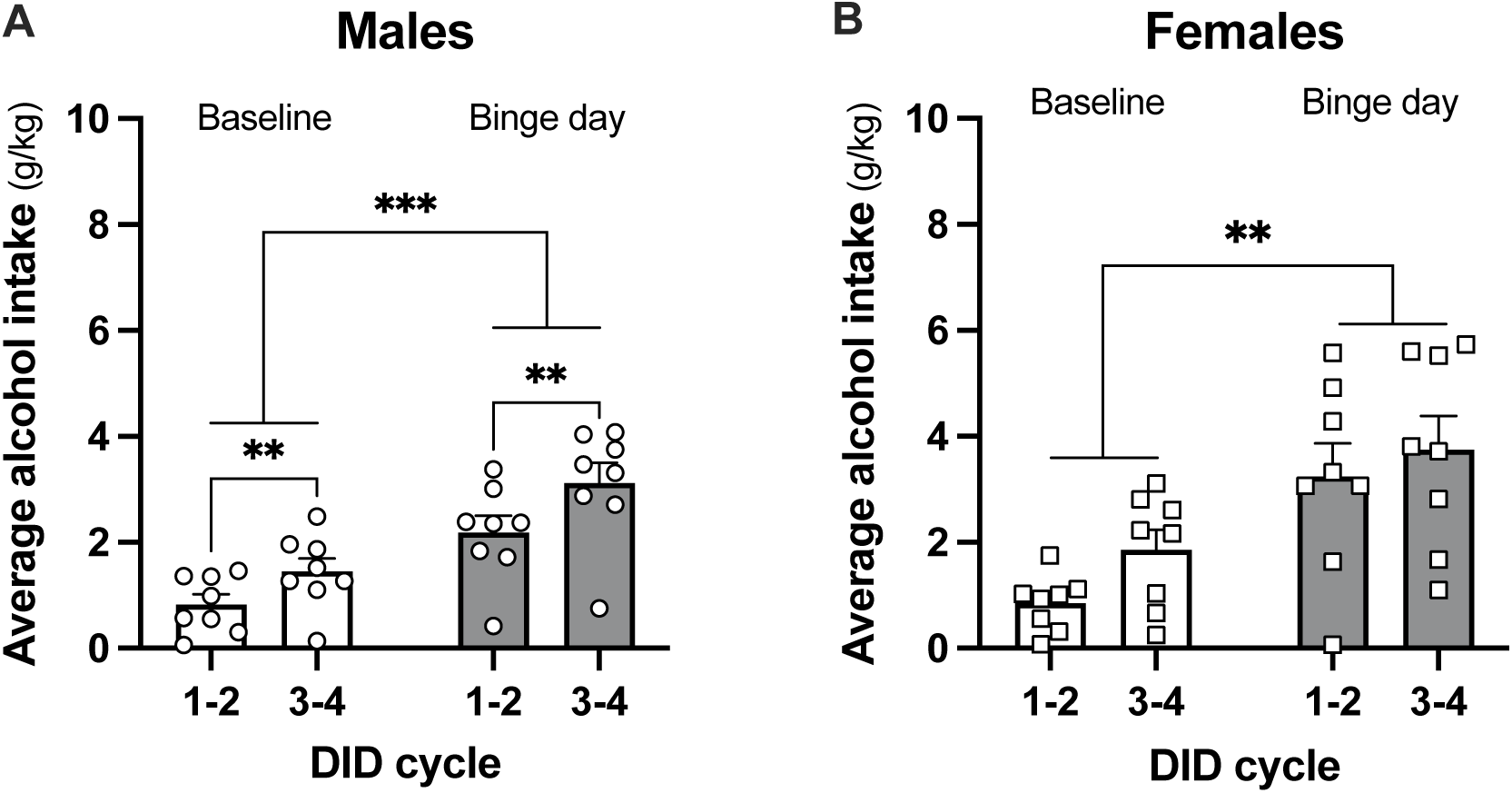
Alcohol intake across adolescent development. **A, B.** Male (**A**) and female (**B**) mice showed a significant increase in the average alcohol intake on binge days compared to baseline days (**, p < 0.01, ***, p < 0.001, two-way ANOVA, main effect of access). Male mice showed an increased average intake during the DID cycles 3 and 4 compared to the average intake in cycles 1 and 2 (**, p < 0.01, two-way ANOVA, main effect of DID cycle). Data are presented as mean values ± SEM with individual values shown in circles (males) or squares (females); *p σ; 0.05 = significant; ns = not significant.

**Supplemental Figure 2.**
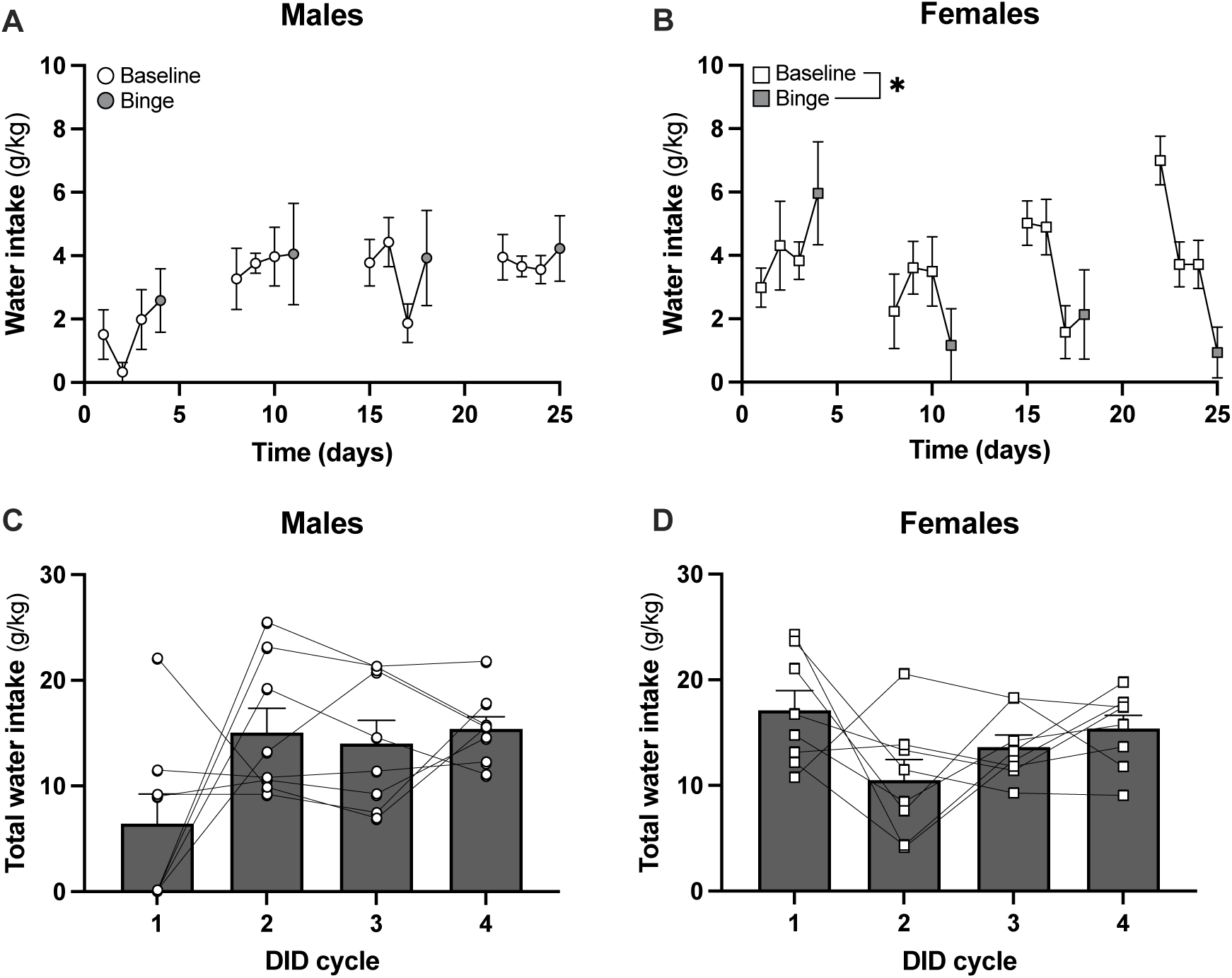
Water intake of adolescent male and female mice. **A.** Male mice did not show consumption difference between baseline and binge days (two-way ANOVA, p > 0.05). **B.** Female mice showed a significant decrease in water intake in the binge session compared to the average of the baseline intake (*, p < 0.05, main effect of access, two-way ANOVA). **C, D.** Water intake in male and female mice showed no differences across the DID cycle (repeated measures one-way ANOVA, p > 0.05). Data are presented as mean values ± SEM with individual values shown in circles (males) or squares (females); *p σ; 0.05 = significant; ns = not significant.

**Supplementary Fig 3.**
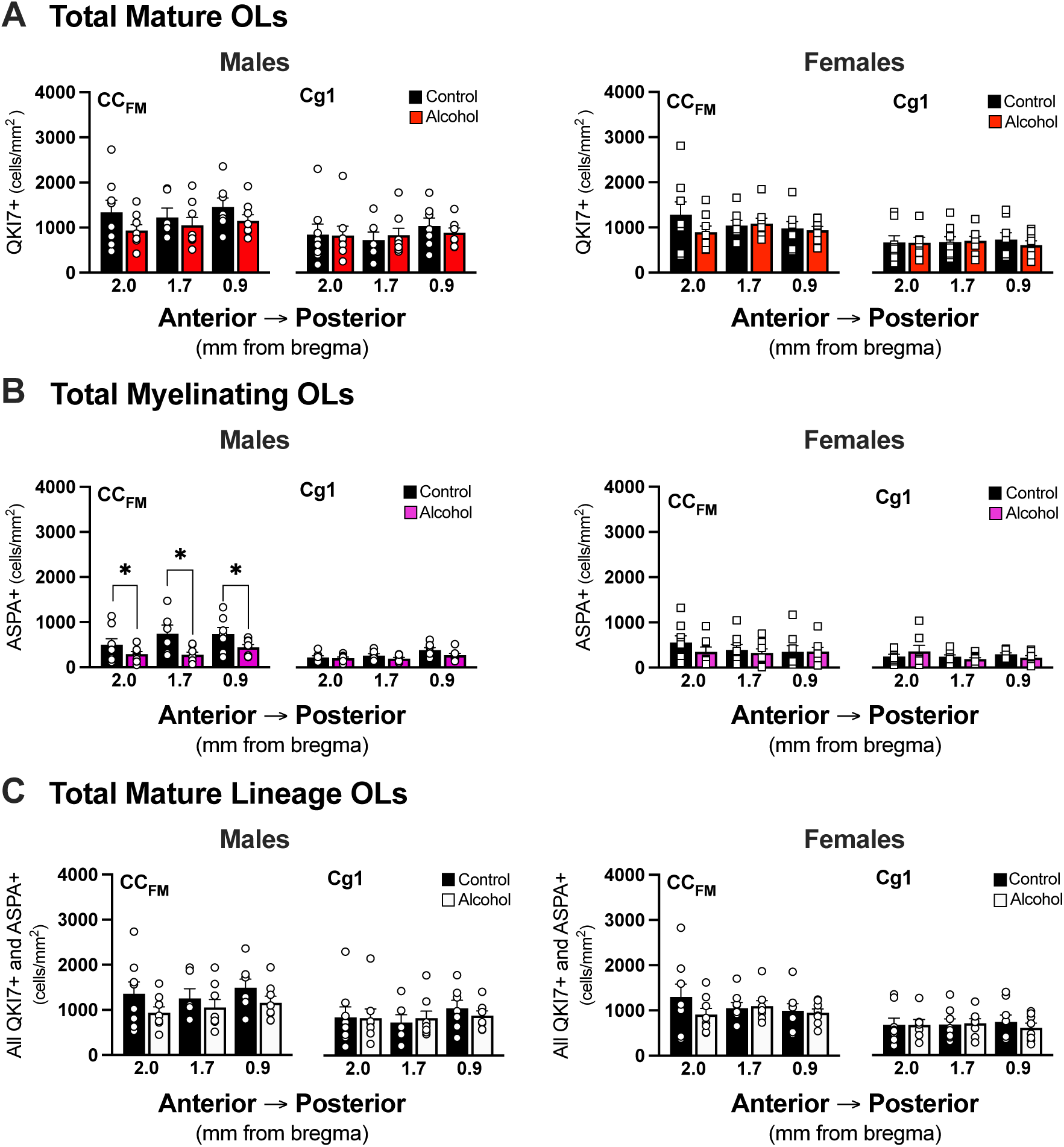
Alcohol decreased the total population density of myelinating OLs with no change in the total population of mature OLs in the CC_FM_ region of male mice. **A.** No differences in the total density of mature (QKI-7+) OLs were found in the CC_FM_ or Cg1 regions of males or females (two-way ANOVA, p > 0.05). **B**. Alcohol decreased the density of total myelinating OLs in the CC_FM_ region of males (*, ps<0.05, main effect of treatment, two-way ANOVA). No differences were found in the CC_FM_ of females or in the Cg1 of males or females (two-way ANOVA, p > 0.05). **C.** No change with alcohol found in the total population of mature OLs in the CC_FM_ or Cg1 regions of males or females (two-way ANOVA, p > 0.05). Data are presented as mean values ± SEM with individual values shown in circles (males) or squares (females); *p σ; 0.05 = significant; ns = not significant.

## References

Acs P, Kipp M, Norkute A, Johann S, Clarner T, Braun A, Berente Z, Komoly S, Beyer C (2009) 17β-estradiol and progesterone prevent cuprizone provoked demyelination of corpus callosum in male mice. Glia 57:807–814.

Addolorato G et al. (2018) Binge Drinking among adolescents is related to the development of Alcohol Use Disorders: results from a Cross-Sectional Study. Sci Rep 8:1–9.

Alfonso-Loeches S, Pascual M, Gómez-Pinedo U, Pascual-Lucas M, Renau-Piqueras J, Guerri C (2012) Toll-like receptor 4 participates in the myelin disruptions associated with chronic alcohol abuse. Glia 60:948–964.

Ambrosius W, Michalak S, Kozubski W, Kalinowska A (2020) Myelin Oligodendrocyte Glycoprotein Antibody-Associated Disease: Current Insights into the Disease Pathophysiology, Diagnosis and Management. Int J Mol Sci 22:100.

American Psychiatric Association (2013) Diagnostic and Statistical Manual of Mental Disorders, 5th ed. Arlinton, VA.

Arancibia-Cárcamo IL, Ford MC, Cossell L, Ishida K, Tohyama K, Attwell D (2017) Node of Ranvier length as a potential regulator of myelinated axon conduction speed. eLife 6:e23329.

Babbs CF, Shi R (2013) Subtle Paranodal Injury Slows Impulse Conduction in a Mathematical Model of Myelinated Axons Hamblin M, ed. PLoS ONE 8:e67767.

Barkley-Levenson AM, Crabbe JC (2012) Ethanol Drinking Microstructure of a High Drinking in the Dark Selected Mouse Line. Alcohol Clin Exp Res 36:1330–1339.

Baslow MH, Suckow RF, Hungund BL (2000) Effects of ethanol and of alcohol dehydrogenase inhibitors on the reduction of N-acetylaspartate levels of brain in mice in vivo: a search for substances that may have therapeutic value in the treatment of Canavan disease. J Inherit Metab Dis 23:684–692.

Baslow MH, Suckow RF, Sapirstein V, Hungund BL (1999) Expression of Aspartoacylase Activity in Cultured Rat Macroglial Cells Is Limited to Oligodendrocytes. J Mol Neurosci 13:47–54.

Baxi EG, DeBruin J, Jin J, Strasburger HJ, Smith MD, Orthmann-Murphy JL, Schott JT, Fairchild AN, Bergles DE, Calabresi PA (2017) Lineage tracing reveals dynamic changes in oligodendrocyte precursor cells following cuprizone-induced demyelination. Glia 65:2087–2098.

Bazzi SA, Maguire C, Mayfield RD, Melamed E (2025) Alcohol induces concentration-dependent transcriptomic changes in oligodendrocytes. Addict Biol 30:e70012.

Bell RL, Rodd ZA, Sable HJK, Schultz JA, Hsu CC, Lumeng L, Murphy JM, McBride WJ (2006) Daily patterns of ethanol drinking in peri-adolescent and adult alcohol-preferring (P) rats. Pharmacol Biochem Behav 83:35–46.

Bin JM, Harris SN, Kennedy TE (2016) The oligodendrocyte-specific antibody ‘CC1’ binds Quaking 7. J Neurochem 139:181–186.

Bowen MT, George O, Muskiewicz DE, Hall FS (2022) Factors contributing to the escalation of alcohol consumption. Neurosci Biobehav Rev 132:730–756.

Cerghet M, Skoff RP, Bessert D, Zhang Z, Mullins C, Ghandour MS (2006) Proliferation and Death of Oligodendrocytes and Myelin Proteins Are Differentially Regulated in Male and Female Rodents. J Neurosci 26:1439–1447.

Chapman TW, Kamen Y, Piedra ET, Hill RA (2024) Oligodendrocyte Maturation Alters the Cell Death Mechanisms That Cause Demyelination. J Neurosci 44:e1794232024.

Chapman TW, Olveda GE, Bame X, Pereira E, Hill RA (2023) Oligodendrocyte death initiates synchronous remyelination to restore cortical myelin patterns in mice. Nat Neurosci 26:555–569.

Chou SP, Pickering RP (1992) Early onset of drinking as a risk factor for lifetime alcohol-related problems. Br J Addict 87:1199–1204.

Clements CS, Reid HH, Beddoe T, Tynan FE, Perugini MA, Johns TG, Bernard CCA, Rossjohn J (2003) The crystal structure of myelin oligodendrocyte glycoprotein, a key autoantigen in multiple sclerosis. Proc Natl Acad Sci 100:11059–11064.

Crabbe JC, Metten P, Rhodes JS, Yu C-H, Brown LL, Phillips TJ, Finn DA (2009) A Line of Mice Selected for High Blood Ethanol Concentrations Shows Drinking in the Dark to Intoxication. Biol Psychiatry 65:662–670.

Creeley CE, Dikranian KT, Johnson SA, Farber NB, Olney JW (2013) Alcohol-induced apoptosis of oligodendrocytes in the fetal macaque brain. Acta Neuropathol Commun 1:23.

Crews FT, Vetreno RP, Broadwater MA, Robinson DL (2016) Adolescent Alcohol Exposure Persistently Impacts Adult Neurobiology and Behavior. Pharmacol Rev 68:1074–1109.

Darbinian N, Darbinyan A, Merabova N, Bajwa A, Tatevosian G, Martirosyan D, Zhao H, Selzer ME, Goetzl L (2021) Ethanol-mediated alterations in oligodendrocyte differentiation in the developing brain. Neurobiol Dis 148:105181.

Dhaher R, McConnell KK, Rodd ZA, McBride WJ, Bell RL (2012) Daily patterns of ethanol drinking in adolescent and adult, male and female, high alcohol drinking (HAD) replicate lines of rats. Pharmacol Biochem Behav 102:540–548.

Dimou L, Simon C, Kirchhoff F, Takebayashi H, Gotz M (2008) Progeny of Olig2-Expressing Progenitors in the Gray and White Matter of the Adult Mouse Cerebral Cortex. J Neurosci 28:10434–10442.

Drzewiecki CM, Willing J, Juraska JM (2020) Influences of age and pubertal status on number and intensity of perineuronal nets in the rat medial prefrontal cortex. Brain Struct Funct 225:2495–2507.

Edwards CM, Xu Z, Winder DG (2025) Adolescent onset of volitional ethanol intake normalizes sex differences observed with adult-onset ethanol intake and negative affective behaviors during protracted forced abstinence. Psychopharmacology (Berl) Available at: https://link.springer.com/10.1007/s00213-025-06925-5 [Accessed January 13, 2026].

Elsayed NM, Kim MJ, Fields KM, Olvera RL, Hariri AR, Williamson DE (2018) Trajectories of Alcohol Initiation and Use During Adolescence: The Role of Stress and Amygdala Reactivity. J Am Acad Child Adolesc Psychiatry 57:550–560.

Flores-Bonilla A, De Oliveira B, Silva-Gotay A, Lucier KW, Richardson HN (2021) Shortening time for access to alcohol drives up front-loading behavior, bringing consumption in male rats to the level of females. Biol Sex Differ 12:51.

Flores-Bonilla A, Richardson HN (2020) Sex Differences in the Neurobiology of Alcohol Use Disorder. Alcohol Res Curr Rev 40:03.

Francis JS, Strande L, Markov V, Leone P (2012) Aspartoacylase Supports Oxidative Energy Metabolism during Myelination. J Cereb Blood Flow Metab 32:1725–1736.

Francis JS, Wojtas I, Markov V, Gray SJ, McCown TJ, Samulski RJ, Bilaniuk LT, Wang D-J, De Vivo DC, Janson CG, Leone P (2016) N-acetylaspartate supports the energetic demands of developmental myelination via oligodendroglial aspartoacylase. Neurobiol Dis 96:323–334.

Gilpin NW, Karanikas CA, Richardson HN (2012) Adolescent Binge Drinking Leads to Changes in Alcohol Drinking, Anxiety, and Amygdalar Corticotropin Releasing Factor Cells in Adulthood in Male Rats Aleman A, ed. PLoS ONE 7:e31466.

Gogtay N, Giedd JN, Lusk L, Hayashi KM, Greenstein D, Vaituzis AC, Nugent TF, Herman DH, Clasen LS, Toga AW, Rapoport JL, Thompson PM (2004) Dynamic mapping of human cortical development during childhood through early adulthood. Proc Natl Acad Sci U S A 101:8174–8179.

Grønbæk-Thygesen M, Hartmann-Petersen R (2024) Cellular and molecular mechanisms of aspartoacylase and its role in Canavan disease. Cell Biosci 14:45.

Guo F, Zhang Y-F, Liu K, Huang X, Li R-X, Wang S-Y, Wang F, Xiao L, Mei F, Li T (2021) Chronic Exposure to Alcohol Inhibits New Myelin Generation in Adult Mouse Brain. Front Cell Neurosci 15:732602.

Hershfield JR, Madhavarao CN, Moffett JR, Benjamins JA, Garbern JY, Namboodiri A, Hershfield JR, Madhavarao CN, Moffett JR, Benjamins JA, Garbern JY, Namboodiri A (2006) Aspartoacylase is a regulated nuclear-cytoplasmic enzyme. FASEB J 20:2139–2141.

Hill RA, Damisah EC, Chen F, Kwan AC, Grutzendler J (2017) Targeted two-photon chemical apoptotic ablation of defined cell types in vivo. Nat Commun 8:15837.

Hill RA, Patel KD, Goncalves CM, Grutzendler J, Nishiyama A (2014) Modulation of oligodendrocyte generation during a critical temporal window aver NG2 cell division. Nat Neurosci 17:1518–1527.

Huang H, He W, Tang T, Qiu M (2023) Immunological Markers for Central Nervous System Glia. Neurosci Bull 39:379–392.

Jacobus J, McQueeny T, Bava S, Schweinsburg BC, Frank LR, Yang TT, Tapert SF (2009) White matter integrity in adolescents with histories of marijuana use and binge drinking. Neurotoxicol Teratol 31:349–355.

Johns TG, Bernard CCA (1999) The Structure and Function of Myelin Oligodendrocyte Glycoprotein. J Neurochem 72:1–9.

Kapuscinski J (1995) DAPI: a DNA-Specific Fluorescent Probe. Biotech Histochem 70:220–233.

Kirmani BF, Jacobowitz DM, Namboodiri MAA (2003) Developmental increase of aspartoacylase in oligodendrocytes parallels CNS myelination. Dev Brain Res 140:105–115.

Lancaster FE, Brown TD, Coker KL, Elliott JA, Wren SB (1996) Sex Differences in Alcohol Preference and Drinking Patterns Emerge during the Early Postpubertal Period in Sprague-Dawley Rats. Alcohol Clin Exp Res 20:1043–1049.

Lee KM, Coehlo MA, Solton NR, Szumlinski KK (2017) Negative Affect and Excessive Alcohol Intake Incubate during Protracted Withdrawal from Binge-Drinking in Adolescent, But Not Adult, Mice. Front Psychol 8:1128.

Lees B, Meredith LR, Kirkland AE, Bryant BE, Squeglia LM (2020) Effect of alcohol use on the adolescent brain and behavior. Pharmacol Biochem Behav 192:172906.

Lehmann ML, Weigel TK, Elkahloun AG, Herkenham M (2017) Chronic social defeat reduces myelination in the mouse medial prefrontal cortex. Sci Rep 7:46548.

Li W, Penderis J, Zhao C, Schumacher M, Franklin R (2006) Females remyelinate more efficiently than males following demyelination in the aged but not young adult CNS. Exp Neurol 202:250–254.

Linsenbardt DN, Timme NM, Lapish CC (2019) Encoding of the Intent to Drink Alcohol by the Prefrontal Cortex Is Blunted in Rats with a Family History of Excessive Drinking. eneuro 6:ENEURO.0489-18.2019.

Liran M, Fischer I, Elboim M, Rahamim N, Gordon T, Urshansky N, Assaf Y, Barak B, Barak S (2025) Long-Term Excessive Alcohol Consumption Enhances Myelination in the Mouse Nucleus Accumbens. J Neurosci 45:e0280242025.

Madhavarao CN, Hammer JA, Quarles RH, Namboodiri MAA (2002) A radiometric assay for aspartoacylase activity in cultured oligodendrocytes. Anal Biochem 308:314–319.

Madhavarao CN, Moffett JR, Moore RA, Viola RE, Namboodiri MAA, Jacobowitz DM (2004) Immunohistochemical localization of aspartoacylase in the rat central nervous system. J Comp Neurol 472:318–329.

Makinodan M, Yamauchi T, Tatsumi K, Okuda H, Takeda T, Kiuchi K, Sadamatsu M, Wanaka A, Kishimoto T (2009) Demyelination in the juvenile period, but not in adulthood, leads to long-lasting cognitive impairment and deficient social interaction in mice. Prog Neuropsychopharmacol Biol Psychiatry 33:978–985.

Mason JL, Jones JJ, Taniike M, Morell P, Suzuki K, Matsushima GK (2000) Mature oligodendrocyte apoptosis precedes IGF-1 production and oligodendrocyte progenitor accumulation and differentiation during demyelination/remyelination. J Neurosci Res 61:251–262.

Mattan NS, Ghiani CA, Lloyd M, Matalon R, Bok D, Casaccia P, De Vellis J (2010) Aspartoacylase deficiency affects early postnatal development of oligodendrocytes and myelination. Neurobiol Dis 40:432–443.

May AC, Stephens LH, Kraybill EP, Meyerhoff DJ, Durazzo TC (2025) Frontal Brain N-Acetylaspartate at Treatment Entry is Related to Future World Health Organization Risk Drinking Levels in Individuals With Alcohol Use Disorder. J Stud Alcohol Drugs 86:416–423.

McCarty CA, Ebel BE, Garrison MM, DiGiuseppe DL, Christakis DA, Rivara FP (2004) Continuity of Binge and Harmful Drinking From Late Adolescence to Early Adulthood. Pediatrics 114:714–719.

McDougall S, Vargas Riad W, Silva-Gotay A, Tavares ER, Harpalani D, Li G-L, Richardson HN (2018) Myelination of Axons Corresponds with Faster Transmission Speed in the Prefrontal Cortex of Developing Male Rats. eneuro 5:ENEURO.0203-18.2018.

Morris VL, Owens MM, Syan SK, Petker TD, Sweet LH, Oshri A, MacKillop J, Amlung M (2019) Associations Between Drinking and Cortical Thickness in Younger Adult Drinkers: Findings From the Human Connectome Project. Alcohol Clin Exp Res 43:1918–1927.

National Institute on Alcohol Abuse and Alcoholism (2004) NIAAA Council Approves Definition of Binge Drinking. NIAAA Newsl 3:3.

Newville J, Valenzuela CF, Li L, Jantzie LL, Cunningham LA (2017) Acute oligodendrocyte loss with persistent white matter injury in a third trimester equivalent mouse model of fetal alcohol spectrum disorder. Glia 65:1317–1332.

Pan S, Mayoral SR, Choi HS, Chan JR, Kheirbek MA (2020) Preservation of a remote fear memory requires new myelin formation. Nat Neurosci 23:487–499.

Pandey SC, Sakharkar AJ, Tang L, Zhang H (2015) Potential role of adolescent alcohol exposure-induced amygdaloid histone modifications in anxiety and alcohol intake during adulthood. Neurobiol Dis 82:607–619.

Papp-Peka A, Tong M, Kril JJ, De La Monte SM, Sutherland GT (2016) The Differential Effects of Alcohol and Nicotine-Specific Nitrosamine Ketone on White Matter Ultrastructure. Alcohol Alcohol:alcalc;agw067v1.

Pascual M, Pla A, Miñarro J, Guerri C (2014) Neuroimmune Activation and Myelin Changes in Adolescent Rats Exposed to High-Dose Alcohol and Associated Cognitive Dysfunction: A Review with Reference to Human Adolescent Drinking. Alcohol Alcohol 49:187–192.

Peters S, Jolles DJ, Duijvenvoorde ACKV, Crone EA, Peper JS (2015) The link between testosterone and amygdala–orbitofrontal cortex connectivity in adolescent alcohol use. Psychoneuroendocrinology 53:117–126.

Poggi G, Albiez J, Pryce CR (2022) Effects of chronic social stress on oligodendrocyte proliferation-maturation and myelin status in prefrontal cortex and amygdala in adult mice. Neurobiol Stress 18:100451.

Radke AK, Sneddon EA, Monroe SC (2021) Studying Sex Differences in Rodent Models of Addictive Behavior. Curr Protoc 1:e119.

Rhodes JS, Best K, Belknap JK, Finn DA, Crabbe JC (2005a) Evaluation of a simple model of ethanol drinking to intoxication in C57BL/6J mice. Physiol Behav 84:53–63.

Rhodes JS, Best K, Belknap JK, Finn DA, Crabbe JC (2005b) Evaluation of a simple model of ethanol drinking to intoxication in C57BL/6J mice. Physiol Behav 84:53–63.

Rice J, Coutellier L, Weiner JL, Gu C (2019) Region-specific interneuron demyelination and heightened anxiety-like behavior induced by adolescent binge alcohol treatment. Acta Neuropathol Commun 7:173.

Rice J, Gu C (2019) Function and Mechanism of Myelin Regulation in Alcohol Abuse and Alcoholism. BioEssays 41:1–9.

Rivers LE, Young KM, Rizzi M, Jamen F, Psachoulia K, Wade A, Kessaris N, Richardson WD (2008) PDGFRA/NG2 glia generate myelinating oligodendrocytes and piriform projection neurons in adult mice. Nat Neurosci 11:1392–1401.

Sacks JJ, Gonzales KR, Bouchery EE, Tomedi LE, Brewer RD (2015) 2010 National and State Costs of Excessive Alcohol Consumption. Am J Prev Med 49:e73–e79.

Schramm-Sapyta NL, Francis R, MacDonald A, Keistler C, O’Neill L, Kuhn CM (2014) Effect of sex on ethanol consumption and conditioned taste aversion in adolescent and adult rats. Psychopharmacology (Berl) 231:1831–1839.

Scurfield A, Latimer DC (2018) A computational study of the impact of inhomogeneous internodal lengths on conduction velocity in myelinated neurons Thomas J-L, ed. PLOS ONE 13:e0191106.

Seidl AH (2014) Regulation of conduction time along axons. Neuroscience 276:126–134.

Silva-Gotay A, Davis J, Tavares ER, Richardson HN (2021) Alcohol drinking during early adolescence activates microglial cells and increases frontolimbic Interleukin-1 beta and Toll-like receptor 4 gene expression, with heightened sensitivity in male rats compared to females. Neuropharmacology 197:108698.

Simons M, Nave K-A (2016) Oligodendrocytes: Myelination and Axonal Support. Cold Spring Harb Perspect Biol 8:a020479.

Sommer WH, Canals S (2025) Alcohol-Induced Changes in Brain Microstructure: Uncovering Novel Pathophysiological Mechanisms of AUD Using Translational DTI in Humans and Rodents. In: Behavioral Neuroscience of Alcohol Addiction (Sommer WH, Spanagel R, eds), pp 595–617 Current Topics in Behavioral Neurosciences. Cham: Springer Nature Switzerland. Available at: https://link.springer.com/10.1007/7854_2025_585 [Accessed March 28, 2026].

Squeglia LM, Tapert SF, Sullivan EV, Jacobus J, Meloy MJ, Rohlfing T, Pfefferbaum A (2015) Brain Development in Heavy-Drinking Adolescents. Am J Psychiatry 172:531–542.

Stadelmann C, Timmler S, Barrantes-Freer A, Simons M (2019) Myelin in the Central Nervous System: Structure, Function, and Pathology. Physiol Rev 99:1381–1431.

Stahre M, Roeber J, Kanny D, Brewer RD, Zhang X (2014) Contribution of excessive alcohol consumption to deaths and years of potential life lost in the United States. Prev Chronic Dis 11:1–12.

Strong MN, Yoneyama N, Fretwell AM, Snelling C, Tanchuck MA, Finn DA (2010) “Binge” drinking experience in adolescent mice shows sex differences and elevated ethanol intake in adulthood. Horm Behav 58:82–90.

Substance Abuse and Mental Health Services Administration (2025a) Highlights for the 2024 National Survey on Drug Use and Health. HHS Publ No PEP25-07-007 NSDUH Ser H-60 Available at: https://www.samhsa.gov/data/sites/default/files/NSDUH%202024%20Annual%20Release/2024-nsduh-nnr-highlights.pdf.

Substance Abuse and Mental Health Services Administration (2025b) Binge Alcohol Use in Past Month: Among People Aged 12 or Older; by Age Group and Demographic Characteristics, Percentages, 2023 and 2024. NSDUH Detailed Table: 2.28B. Available at: https://www.samhsa.gov/data/sites/default/files/reports/rpt56484/NSDUHDetailedTabs2024/NSDUHDetailedTabs2024/2024-nsduh-detailed-tables-sect2pe.htm#tab2.28b.

Substance Abuse and Mental Health Services Administration (2025c) Key substance use and mental health indicators in the United States: Results from the 2024 National Survey on Drug Use and Health. HHS Publ No PEP25-07-007 NSDUH Ser H-60 Available at: https://www.samhsa.gov/data/sites/default/files/reports/rpt56287/2024-nsduh-annual-national-report.pdf.

Takeda S, Hoshiai R, Tanaka M, Izawa T, Yamate J, Kuramoto T, Kuwamura M (2024) Myelin lesion in the aspartoacylase (*Aspa*) knockout rat, an animal model for Canavan disease. Exp Anim 73:347–356.

Tavares, Silva-Gotay, Riad, Bengston, Richardson (2019) Sex Differences in the Effect of Alcohol Drinking on Myelinated Axons in the Anterior Cingulate Cortex of Adolescent Rats. Brain Sci 9:167.

Taylor LC, Puranam K, Gilmore W, Ting JP-Y, Matsushima GK (2010) 17β-estradiol protects male mice from cuprizone-induced demyelination and oligodendrocyte loss. Neurobiol Dis 39:127–137.

Thiele TE, Crabbe JC, Boehm SL (2014) “Drinking in the Dark” (DID): A Simple Mouse Model of Binge-Like Alcohol Intake. Curr Protoc Neurosci 68:1–12.

Thiele TE, Navarro M (2014) “Drinking in the dark” (DID) procedures: A model of binge-like ethanol drinking in non-dependent mice. Alcohol 48:235–241.

Vargas WM, Bengston L, Gilpin NW, Whitcomb BW, Richardson HN (2014) Alcohol Binge Drinking during Adolescence or Dependence during Adulthood Reduces Prefrontal Myelin in Male Rats. J Neurosci 34:14777–14782.

Viganò F, Möbius W, Götz M, Dimou L (2013) Transplantation reveals regional differences in oligodendrocyte differentiation in the adult brain. Nat Neurosci 16:1370–1372.

Walker BM, Walker JL, Ehlers CL (2008) Dissociable effects of ethanol consumption during the light and dark phase in adolescent and adult Wistar rats. Alcohol 42:83–89.

Wilcox M V., Carlson VCC, Sherazee N, Sprow GM, Bock R, Thiele TE, Lovinger DM, Alvarez VA (2014) Repeated Binge-like ethanol drinking alters ethanol drinking patterns and depresses striatal GABAergic transmission. Neuropsychopharmacology 39:579–594.

Wolstenholme JT, Mahmood T, Harris GM, Abbas S, Miles MF (2017) Intermittent Ethanol during Adolescence Leads to Lasting Behavioral Changes in Adulthood and Alters Gene Expression and Histone Methylation in the PFC. Front Mol Neurosci 10:307.

Young KM, Psachoulia K, Tripathi RB, Dunn S-J, Cossell L, Attwell D, Tohyama K, Richardson WD (2013) Oligodendrocyte Dynamics in the Healthy Adult CNS: Evidence for Myelin Remodeling. Neuron 77:873–885.

Younis RM, Wolstenholme JT, Bagdas D, BeÅnger JC, Miles MF, Damaj MI (2019) Adolescent but not adult ethanol binge drinking modulates ethanol behavioral effects in mice later in life. Pharmacol Biochem Behav 184:172740.

